# Multi-omic analysis of subtype evolution and heterogeneity in high-grade serous ovarian carcinoma

**DOI:** 10.1101/554394

**Authors:** Ludwig Geistlinger, Sehyun Oh, Marcel Ramos, Lucas Schiffer, Rebecca LaRue, Christine Henzler, Sarah Munro, Claire Daughters, Andrew C. Nelson, Boris Winterhoff, Zenas Chang, Shobhana Talukdar, Mihir Shetty, Sally Mullaney, Martin Morgan, Giovanni Parmigiani, Michael Birrer, Li-Xuan Qin, Markus Riester, Timothy K. Starr, Levi Waldron

## Abstract

Multiple studies have identified transcriptome subtypes of high-grade serous ovarian carcinoma (HGSOC), but these have yet to impact clinical practice. Interpretation and translation of HGSOC subtypes are complicated by tumor evolution and polyclonality accompanied by accumulation of somatic aberrations, varying cell type admixtures, and different tissues of origin. The chronology of HGSOC subtype evolution was examined in the context of these factors by a novel integrative analysis of bulk absolute somatic copy number analysis and gene expression in The Cancer Genome Atlas, complemented by single-cell RNA-seq analysis of six independent tumors. The approach was validated by contrast to soft-tissue sarcoma. Genomic lesions associated with HGSOC subtypes tend to be subclonal, implying subtype divergence at later stages of tumor evolution. Subclonality of recurrent HGSOC alterations is particularly evident for proliferative tumors, characterized by extreme genomic instability, absence of immune infiltration, and greater patient age. In contrast, differentiated tumors are characterized by largely intact genome integrity, high immune infiltration, and younger patient age. We propose an alternative model to discrete subtypes of HGSOC, in which tumors develop from an early differentiated spectrum to a late proliferative spectrum, along a timeline characterized by increasing genomic instability and subclonal expansion. The proposed methods provide a new approach to investigating tumor evolution through multi-omic analysis.

**Statement of Significance:** This study proposes a method to infer whether transcriptome-based groupings of tumors differentiate early in carcinogenesis and are therefore potentially appropriate targets for therapy, and demonstrates that this is not the case for high-grade serous ovarian carcinoma (HGSOC). Significant findings for HGSOC include:

- Tumor purity, ploidy, and subclonality can be reliably inferred from different genomic platforms and show marked differences between subtypes
- Recurrent DNA alterations are associated with subtypes and tend to occur more frequently in subclones
- Single-cell sequencing of 42,000 tumor cells reveals widespread heterogeneity in tumor cell type composition that drives bulk subtype calls, but demonstrates a lack of intrinsic subtypes among tumor epithelial cells
- Findings prompt the dismissal of discrete transcriptome subtypes for HGSOC and replacement by a more realistic model of continuous tumor development that includes mixtures of subclones, accumulation of somatic aberrations, infiltration of immune and stromal cells in proportions correlated with tissue of origin and tumor stage, and evolution between properties previously associated with discrete subtypes

## Introduction

High-grade serous ovarian cancer (HGSOC) is the most common histological subtype of ovarian cancer, accounts for 70-80% of ovarian cancer deaths, and is associated with poor prognosis and frequent relapse^1,2^. HGS ovarian cancer is a genomically complex disease that is characterized by ubiquitous TP53 mutations^3^, frequent loss of *RB1, NF1* and *PTEN* by gene breakage events^4,5^, and recurrent high-level copy number amplifications^6^.

Molecular stratification of HGSOC is difficult due to the genomic complexity and extensive heterogeneity of the disease. Clinically relevant genomic stratification is currently restricted to the identification of homologous recombination deficiency (HRD), a condition that is present in roughly half of all HGS ovarian tumors, and that is attributable to germline or somatic BRCA mutations in approximately 20% of HGSOC cases^7,8^.

Several studies also reported molecularly distinct subtypes by clustering tumors together that have similar transcriptome profiles^4,9–12^. The Cancer Genome Atlas (TCGA) project reported four subtypes^4^ and named them based on marker gene expression: *differentiated, immunoreactive, mesenchymal*, and *proliferative*. These subtypes were found associated with several clinical and tumor pathology characteristics^4,11,13^, potentially reflect different tissues of origin^14^, and partly reflect differences in immune cell^15^ and stromal^16^ content. Robustness and clinical utility of the transcriptome subtypes remain controversial^17,18^. Based on a compendium of 15 microarray datasets consisting of ≈1,800 HGSOC tumors, subtype classifiers were not robust to re-fitting in independent datasets and grouped only one third of patients concordantly into four subtypes^13^.

Most HGSOC tumors are polyclonal, meaning that a single tumor is a heterogeneous assembly of distinct cancer genotypes arising from different subclones. Lohr *et al.* estimated that 95% of the TCGA HGSOC tumors are polyclonal, and ≈40% consist of ≥4 subclones^19^. Recent single-cell studies further demonstrated extensive intra-tumoral heterogeneity of HGS ovarian tumors^20–23^, consistent with the notion of an individual tumor being an intermixture of different malignant cell populations. Transcriptome subtypes might provide a useful summary of bulk tumor properties if these subtypes are shared by different intratumoral subclones, representing an intrinsic difference between tumors. However, we hypothesized that subtype assignment via transcriptome clustering is driven by late events in tumor evolution, so that these subtype properties would be subclone-specific. We test this hypothesis first through a novel use of allele-specific copy number of genomic regions identified as subtype markers in bulk tumors. In contrast to phylogenetic analysis based on longitudinal whole-genome sequencing at multiple time points^24,25^, the proposed approach infers tumor evolution from copy number data obtained with genotyping arrays or exome sequencing at a single time point. We follow up with single-cell RNA-seq (scRNA-seq) analysis of six independent patients and investigate whether individual epithelial cancer cells have different well-defined subtypes and whether they correspond to the proposed subtypes of bulk tumors.

## Materials & Methods

Statistical analysis was carried out using R^26^ and Bioconductor^27^. Code is available from GitHub (https://github.com/waldronlab/subtypeHeterogeneity).

### SCNA subtype association

SCNAs detected with GISTIC2^28^ were obtained from the final run of the TCGA Firehose pipeline (2016-01-28). SCNAs were classified depending on their type (deletion / amplification) as either normal (0), loss / gain of a single copy (1), or loss / gain of two or more copies (2). Transcriptome clusters, assigning each tumor to a subtype^4^, were retrieved from the 2016-01-28 Firehose run. Subtype association was tested by *χ*^2^ test with *df* = 6. Multiple testing correction was carried out using an FDR^29^ cutoff of 0.1.

### SCNA subclonality

ABSOLUTE^30^ SCNA calls were obtained from the Pan-Cancer Atlas aneuploidy study^31^. This included per-tumor estimates of purity, ploidy, subclonal genome fraction, number of genome doublings, and segmented absolute CN calls classified as occurring clonally or subclonally. ABSOLUTE calls were managed in the R/Bioconductor data class RaggedExperiment, which facilitated summarization of ABSOLUTE’s subclonality calls in GISTIC2 regions using the qreduceAssay function. A region was called subclonal if overlapped by at least one subclonality call. GISTIC2 peaks were extended by 500 kb up- and downstream to account for uncertainty of the peak calling heuristic.

### Correlation of subtype association with subclonality

Using the *χ*^2^ test statistic as the subtype association score *S*_*A*_ of an alteration (Figure 3A), and the fraction of tumors for which this alteration is subclonal as the subclonality score *S*_*C*_ (Figure 3B), Spearman’s rank correlation was computed to assess the relationship between *S*_*A*_ and *S*_*C*_. Statistical significance of the correlation was assessed using Spearman’s rank correlation test. To account for non-independence of the occurrence of different SCNAs, we also carried out a permutation test, where we permuted the observed *S*_*A*_ values 1000 times and recalculated the correlation with the observed *S*_*C*_ values. The *p*-value was obtained by calculating the fraction of permutations in which the correlation of the permuted setup exceeded the observed correlation.

**Figure 1:**
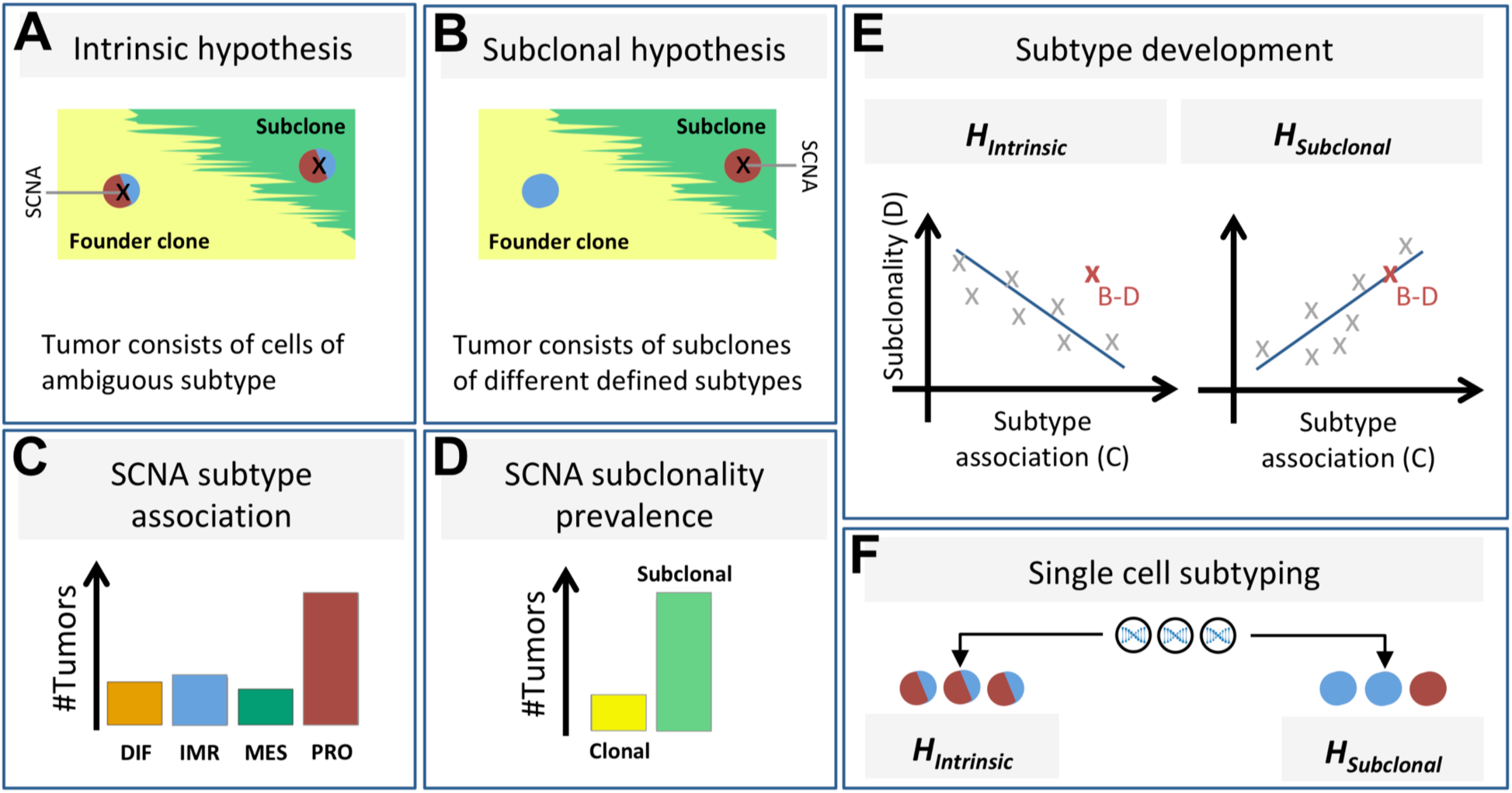
Study setup. Our study aims to distinguish between two possible hypotheses explaining why gene expression-based HGSOC subtypes are ambiguous. The intrinsic hypothesis **(A)** is that tumor cells display ambiguous expression patterns consisting of two or more subtype expression patterns. The subclonal hypothesis **(B)** is that a tumor contains multiple clones, with each clone displaying a consistent, yet distinct subtype expression pattern. To distinguish between these two hypotheses, we analyze recurrent SCNAs across many tumors and determine for each SCNA whether it occurs disproportionately often in tumors of a specific subtype **(C)**, and whether it occurs in the founder clone or a subclone **(D)**. The bar charts in **(C)** and **(D)** show a particular SCNA associated with the proliferative subtype, occurring predominantly subclonally. If the subclonal hypothesis were true, there should be a positive correlation between SCNA subtype association and SCNA subclonality prevalence, while the intrinsic hypothesis predicts a negative correlation **(E)**. For example, the SCNA depicted in **(B-D)** (high subtype association and high subclonality) is more consistent with the subclonal hypothesis than with the intrinsic hypothesis (red X in **E**). However, only a trend across many recurrent SCNAs is considered evidence for either hypothesis. Analysis of single cell gene expression patterns **(F)** should also distinguish between the two hypotheses.

**Figure 2:**
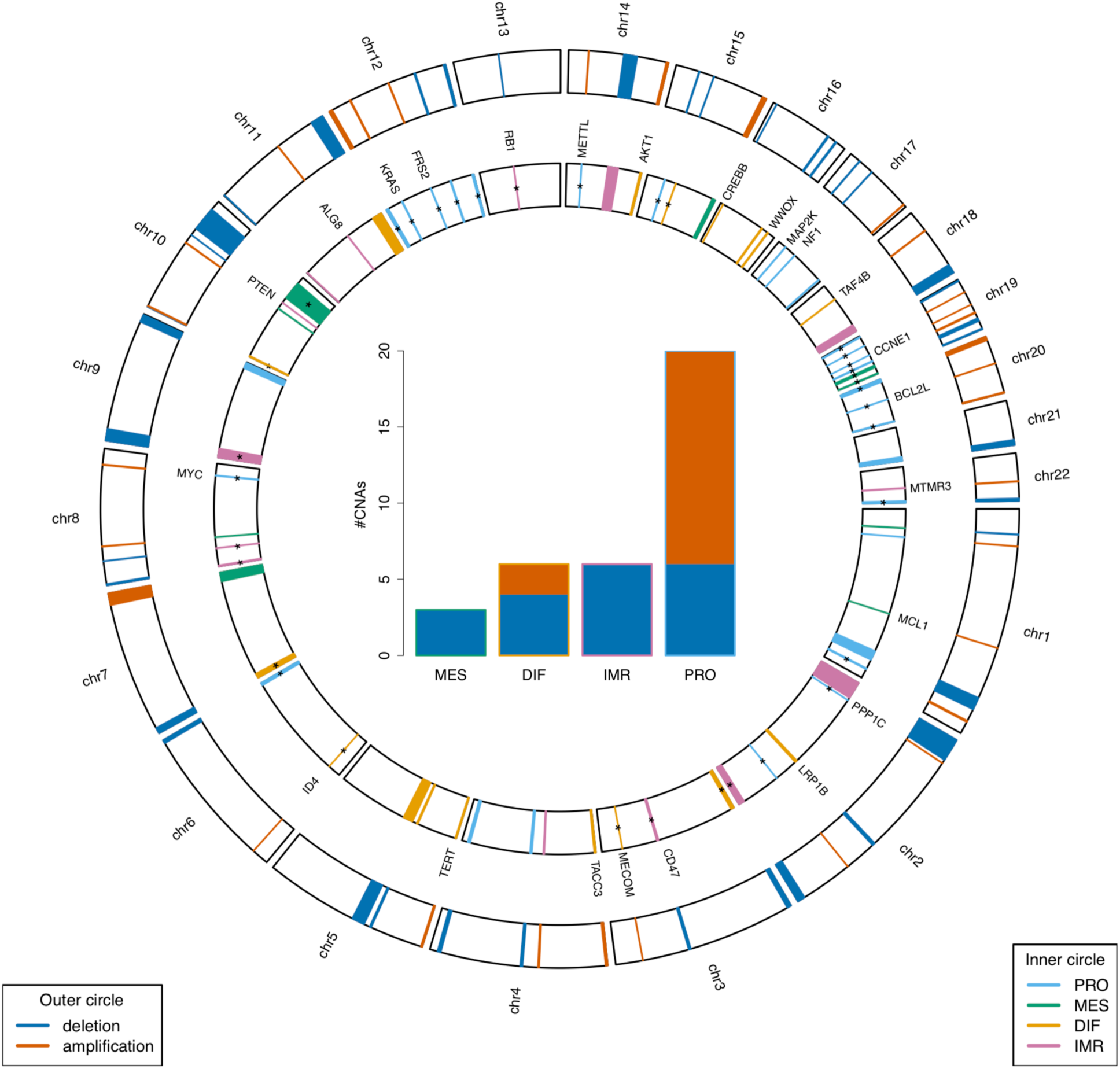
Genomic distribution of subtype-associated SCNAs. The circle on the outside shows the genomic location of focal CN amplifications (red) and deletions (blue) as detected with GISTIC2^28^ in TCGA HGS ovarian tumors. In the inner circle, the detected SCNAs are colored according to subtype association (blue: proliferative, green: mesenchymal, orange: differentiated, violet: immunoreactive). A star indicates significant association (FDR cutoff of 0.1, *χ*^2^ test, Supplementary Figure S2). For example, the *MYC*-containing amplification on chromosome 8 is significantly associated with the proliferative subtype as previously reported^4^. The barplot in the center shows for each subtype (*x*-axis) the number of significantly associated SCNAs (FDR < 0.1, *y*-axis) classified as deletion (blue) or amplification (red). Associations with the proliferative subtype are significantly overrepresented among the subtype-associated regions (20 out of 35, *p* = 0.007, Fisher’s exact test).

**Figure 3:**
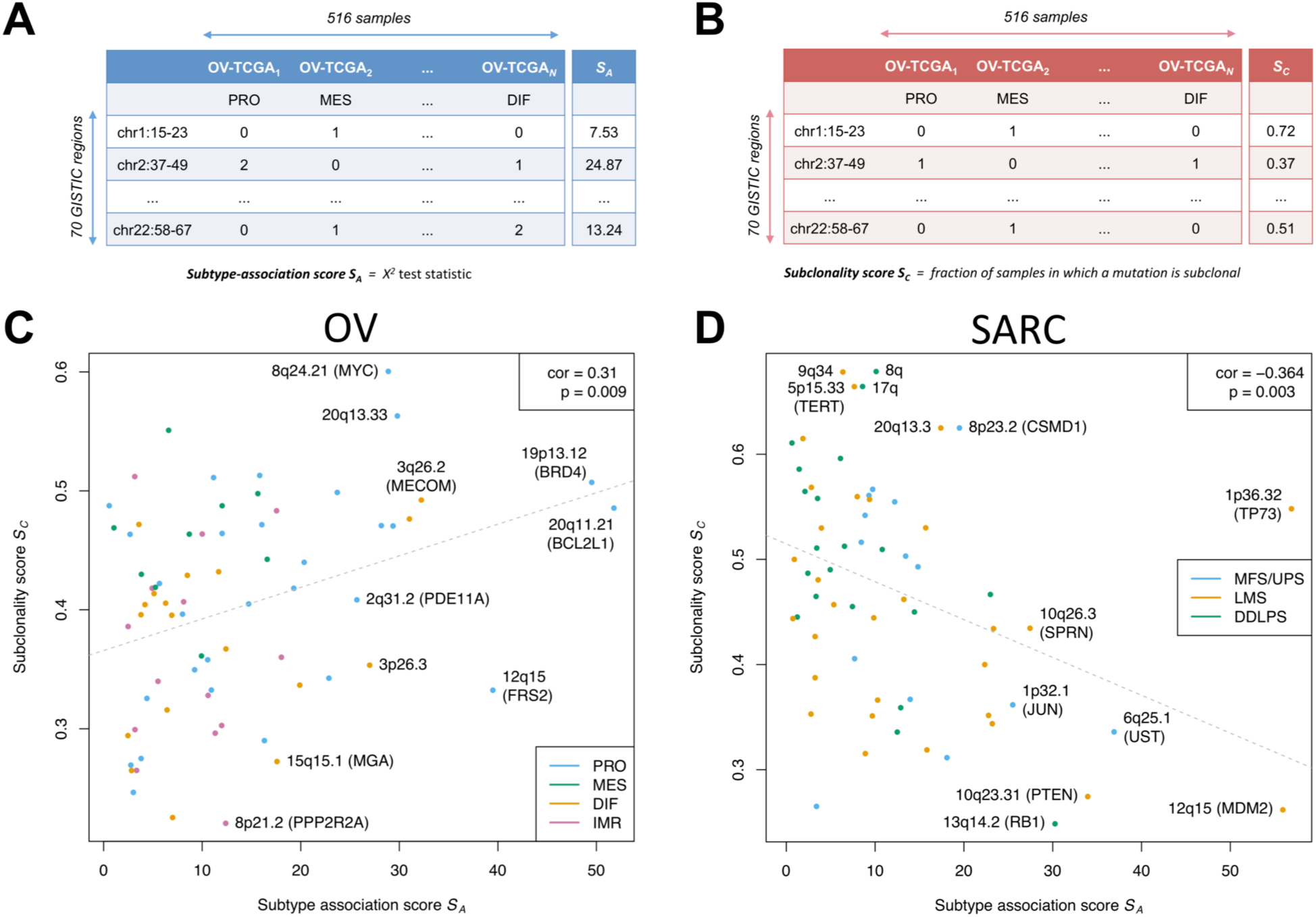
Correlation between subtype association and subclonality. The table in **(A)** illustrates the computation of the subtype-association score *S*_*A*_. Rows of the table correspond to the focal SCNAs detected with GISTIC2^28^. Columns correspond to the 516 TCGA HGS ovarian tumors, each one assigned to one of the 4 subtypes by transcriptome clustering (2nd row). The cells of the table indicate whether the region is of normal state (0), or contains a single (1) or higher (2) copy gain / loss, depending on whether the SCNA is rendered an amplification or deletion. The computation of the subclonality score *S*_*C*_ is analogously illustrated in **(B)**. Here, the table cells indicate whether an SCNA is called as subclonal (1) or not (0) by ABSOLUTE^30^. The scatter plot in **(C)** depicts the correlation between *S*_*A*_ and *S*_*C*_, showing a significant positive correlation of 0.31 with a *p*-value of 0.009 (Spearman’s correlation test). The color of the dots corresponds to subtype association as in Figure 2. Correlation when stratifying tumors by purity is shown in Supplementary Figure S3. Repeating the analysis outlined in **(A-C)** for TCGA soft tissue sarcomas (64 focal SCNAs, 259 samples) results in a significant negative correlation of -0.36 with a *p*-value of 0.003 as depicted in **(D)**.

### Transcriptome subtype classification with the consensusOV classifier

Subtypes were classified using the consensus classifier implemented in the function get.consensus.subtypes of the consensusOV Bioconductor package^13^. The consensus classifier calculates a score (more precisely: a probability in [0,1]) for each subtype and assigns each tumor to the subtype with the highest score. A tumor with a large difference or *margin* between the highest and second highest scores can be considered confidently classifiable, whereas a tumor with two nearly equal scores can be considered of ambiguous subtype.

### Single-cell analysis

Five additional HGSOC specimens were subjected to 10x Genomics single-cell RNA sequencing, producing an average of 235 million reads per patient sample with an average of 11,502 cells/sample and 23,000 reads/cell. Statistical analysis was carried out using a collection of Bioconductor packages for single-cell analysis^32^. Sample collection, experimental procedures, and data processing steps are described in Supplementary Methods S1.

## Results

We previously reported a systematic assessment of four reported HGSOC subtypes with respect to robustness and association to overall survival, and found that most tumors cannot be classified reliably, but that it is possible to predict how reliably each tumor can be classified and that some tumors can be classified with high confidence^13^. Here, we investigate an alternative model of HGSOC development in which ambiguity in tumor classification arises as a consequence of accumulated mutations and intra-tumor heterogeneity (Figure 1). To test this hypothesis, we focus on somatic copy number alterations (SCNAs) given their causal roles in oncogenesis and their potential to discriminate between cancer subtypes^33–35^. The GISTIC2 method detects SCNAs that are more recurrent than expected by chance, in order to distinguish cancer-driving events from random passenger alterations^28^. The method also separates broad *arm-level* events from narrow *focal* events, which often harbor oncogenes and tumor suppressors^36,37^. The ABSOLUTE algorithm^30^ infers tumor purity and ploidy from the analysis of SCNAs, and incorporates this information for quantification of an alteration’s absolute CN per cancer cell. The algorithm also identifies SCNAs not fitting a tumor’s purity and ploidy relationship as a consequence of subclonal evolution.

Focusing on the subtypes proposed by TCGA^4^, we integrate information from 516 TCGA cases GISTIC2 and ABSOLUTE SCNA calls to analyze whether recurrent subtype-associated SCNAs display greater intra-tumor heterogeneity than other alterations. We assess the reliability of these calls by absolute CN analysis on whole-exome sequencing data, and complement results with single-cell subtype classification on approximately 42,000 cells from six independent tumors.

### Tumor purity, ploidy, and subclonality can be reliably inferred from different genomic platforms and show marked differences between subtypes

Previous studies reported specific clinical and tumor pathology characteristics of the four subtypes^4,13^. Using previously published^31^, curated allele-specific copy number estimates obtained by ABSOLUTE analysis, we observed significant differences in tumor purity (defined as the proportion of malignant cells) between subtypes (Supplementary Figure S1a) in agreement with previous reports^38^. Tumors of differentiated subtype were characterized by high purity, but lower ploidy and subclonality than the other three subtypes (Supplementary Figure S1b,c), consistent with a lower number of genome doublings (Supplementary Figure S1d).

To establish the reliability of inferring SCNA subclonality with ABSOLUTE from SNP-array data, we investigated whether results are consistent when using whole-exome sequencing data instead. In a separate benchmarking study^39^, we compared results from absolute copy number analysis of genotyping arrays with matched normal samples (the approach used here) to allele-specific copy number analysis of exome sequencing with and without matched normal samples, using the PureCN R/Bioconductor package^40^. There^39^ we report that per-tumor estimates of purity and ploidy were in good agreement between experimental platforms and computational methods (Pearson correlation of 0.77 for purity and 0.74 for ploidy). This also applied to individual CN calls in GISTIC2 regions, where we found a median concordance of 87.7% of tumors with identical CN state for one GISTIC2 region at a time.

### Recurrent DNA alterations are associated with subtypes and tend to occur more frequently in subclones

We analyzed recurrent focal SCNAs as identified with GISTIC2 in TCGA HGSOC tumors for association with subtypes (Figure 2). We tested all 70 SCNAs identified by GISTIC2, comprising 31 amplifications and 39 deletions (Figure 2, outer ring). Nominal *p*-values showed a concentration near zero (Supplementary Figure S2a), corresponding to 35 alterations being significantly associated with subtypes (FDR < 0.1, Figure 2, inner ring). Associations with the proliferative subtype were significantly overrepresented (20 of 35, *p* = 0.007, Fisher’s exact test, Figure 2, barplot). Strongest subtype association was observed for the *FRS2* and *BLC2L1* amplifications (Figure 2, gene names).

We test the hypothesis that reported HGSOC subtypes differentiate late in tumorigenesis by assessing the correlation between subtype association and subclonality of recurrent SCNAs (Figure 1A-E). Subtype association of an alteration is calculated via a score *S*_*A*_, corresponding to the *χ*^2^ test statistic (Figure 3A). Subclonality prevalence of an alteration is calculated via a score *S*_*C*_, defined as the fraction of samples for which this alteration is subclonal (Figure 3B).

Under the null hypothesis that subtype-associated alterations occur no earlier or later than other alterations, Spearman correlation *ρ* between *S*_*A*_ and *S*_*C*_ would be expected to be zero:

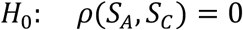

Rejection of *H*_0_ has clear interpretation: if subtype-associated SCNAs tend to be subclonal, i.e.

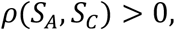

this suggests that the subtypes are late events in tumor evolution. If subtype-associated alterations tend not to be subclonal, i.e.

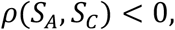

this instead suggests that subtypes are early events, consistent with these being “intrinsic” subtypes.

We obtained a significant positive correlation between subtype association and subclonality prevalence of the 70 SCNAs (Figure 3C). To account for non-independence of the occurrence of different SCNAs, we carried out a permutation test (*p* = 0.006). When stratifying tumors by purity, since very high or low purity tumors present challenges for allele-specific copy number analysis, the correlation was positive in all strata and did not significantly differ between strata (Supplementary Figure S3). Highest subtype association and subclonality were observed for amplifications, and repeatedly displayed increased alteration frequency for the proliferative subtype (including amplification of *BRD4* and the telomeric 20q13.33, Supplementary Figures S4 and S5). A notable exception was the highly subclonal *MYC* amplification, which displayed decreased alteration frequency for proliferative tumors as previously reported^4^. Predominantly clonal alterations were enriched for deletions (9 of 10 regions with *S*_*C*_ < 0.3) with moderate subtype association (including loss of *PPP2R2A* and *MGA*, Supplementary Figure S4). Consistent with the reported frequent loss of *PTEN, RB1*, and *NF1* in HGSOC^4,8^, those alterations occurred predominantly clonally and largely irrespective of subtype classification (Supplementary Figure S6).

As an opposing example, we repeated the analysis for adult soft tissue sarcoma (STS), which like HGSOC is characterized by high SCNA frequency; however, unlike HGSOC, TCGA ST sarcomas are comprised of sarcoma types from different anatomical sites that are characterized by distinct genomic alterations, consistent with them being true intrinsic subtypes^41^. In contrast with HGSOC, association with the STS type-dominated transcriptome clusters was negatively correlated with subclonality prevalence (Figure 3D, Supplementary Results 2.1), consistent with STS being characterized by early / intrinsic type-specific events^41,42^.

### Single-cell sequencing of 42,000 tumor cells reveals widespread heterogeneity in tumor composition and intrinsic ambiguity in subtype assignment

We assessed whether subtype classification ambiguity exists at the level of single cells, or arises from individual epithelial cells of the same tumor being classified as different subtypes (Figure 1F). We therefore analyzed data from Fluidigm deep sequencing of 66 cells from a previously published tumor^20^, and 10x Genomics shallow sequencing of 42,000 cells from five additional tumors. We applied the consensusOV subtype classifier, previously trained on tumors concordantly classified by three major subtype classifiers across 15 microarray datasets^13^, to single cells in conjunction with cell type classification. The consensusOV classifier also displayed high concordance when classifying TCGA tumors assayed by bulk RNA-seq and microarray (Supplementary Figure S7A).

From application of the consensus classifier to bulk RNA-seq and Fluidigm single-cell RNA-seq of 66 cells from a previously published tumor^20^, we observed higher subtype ambiguity for single cells when compared to the bulk tumor (Supplementary Results S2.2). The observed ambiguity in classification of single cells could not be attributed to the low coverage of scRNA-seq data^43^, as downsampling of bulk RNA-seq data to match the coverage of the scRNA-seq data still allowed for a more confident subtype assignment (Supplementary Figure S7B). This also demonstrated new challenges for subtype classification of single cells, as many cells substantially expressed only small parts of the subtype signatures derived from bulk tumors (Supplementary Figure S7C).

To investigate whether these observations hold when analyzing the full complement of tumor cells on a larger scale, we employed 10x Genomics scRNA-seq for five additional tumors. The tumors were clinically classified as stage III-IV, grade 3, and displayed different responses to chemotherapy (Supplementary Table S1). After quality control, the number of cells analyzed for each tumor ranged from 3,630 to 13,747 cells, comprising a total of 42,253 individual tumor cells.

Cell type annotation via marker gene expression and transcriptome similarity to pure cell types (Supplementary Figures S9-S11) consistently returned 5 major cell type populations: epithelial, endothelial, lymphocyte, myeloid, and stromal (Figure 4B,D). However, the respective proportion of each cell type differed significantly between tumors (Figure 4D, proportion test, *df* = 4, *p* < 2.2×10^−16^). Noting that a dedicated single-cell classifier should simultaneously classify cell type and subtype of cancerous epithelial cells, we compared cell type assignments with subtype calls of the `consensusOV` classifier. This demonstrated that certain subtype calls tend to coincide with specific cell types (Figure 4A,C,E). This was evident for immunoreactive calls, found on myeloid innate immune cells (79.5%), and mesenchymal calls that coincided with stromal cells (93.2%, Figure 4E). Epithelial cells were classified as differentiated (84.8%) and to a lesser extent proliferative (13.7%). The cellular tumor composition was reflected in the subtype classification of bulk RNA-seq data for the 5 tumors (Figure 4F, upper panel). Tumor T59 was classified as immunoreactive, consistent with large immune cell populations (Figure 4A,B). Tumor T77 was classified borderline DIF/PRO in agreement with two large DIF/PRO epithelial cell populations, and Tumor T90 was classified as proliferative. Tumors T76 and T89 were confidently classified as MES and IMR, recapturing large stromal and immune cell populations. Comparing the margin scores (differences between the probabilities of most-probable and second most-probable subtype, representing certainty of classification) of tumor epithelial cells with the bulk tumor (Figure 4F, lower panel) demonstrated that classification of individual cells was at least as uncertain as for the bulk, consistent with the observations made for the first tumor (Supplementary Figure S7B). Subtype calls on epithelial cells of Tumors T59 and T77 displayed transitional patterns between differentiated and proliferative (Supplementary Figure S12), similar to the transitional pattern between differentiated and immunoreactive for the first tumor (Supplementary Figure S7C).

**Figure 4:**
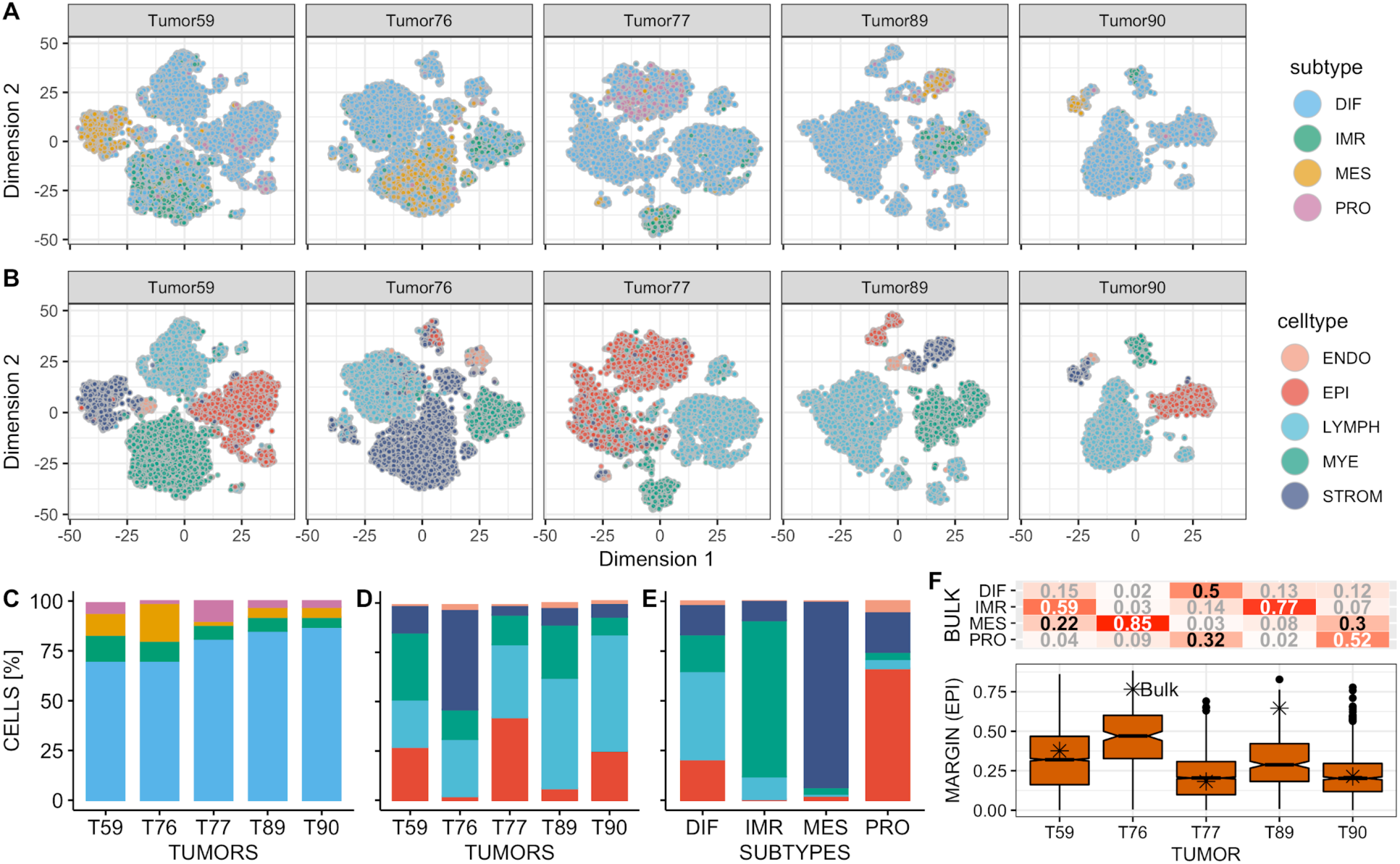
Composition of HGS ovarian tumors at single-cell level. **(A)** and **(B)** show subtype and cell type assignments for 42,253 individual tumor cells of 5 tumors. **(C)** and **(D)** show the proportion of subtype and cell type calls for all five tumors. **(E)** shows the percentage of cells of a specific cell type coinciding with the 4 subtype calls across the 5 tumors. **(F)** shows subtype classification probabilities for bulk RNA-seq data of the 5 tumors (upper panel), and margin score distribution for the epithelial cells of the respective tumor (lower panel). The margin score of the bulk tumor is indicated by an asterisk (higher margin values correspond to more confident subtype assignment).

As epithelial cells were classified as either differentiated or proliferative, we next investigated which factors drive subtype classification in these two groups of cells by analyzing them for differences in copy number profile, cell cycle activity, and expression of bulk subtype markers (Figure 5). Copy number profiles inferred from the scRNA-seq data using inferCNV^44^ demonstrated genetic heterogeneity between tumors, each with an individual mutational landscape shaped by recurrent bulk SCNAs that were also present on single-cell level (Figure 5A and Supplementary Figures S13-S15). The small epithelial cell populations of Tumors T76 and T89 had relatively few copy number alterations: T76 was dominated by two arm-level amplifications on chromosome 8 and 20; T89 carried a *PAX8* amplification that was not seen in the other tumors. The vast majority of epithelial cells of both tumors were confidently classified as differentiated (mean margin of 0.47 and 0.31, respectively). Tumor T59 showed characteristic indicators of homologous recombination deficiency, including strong amplifications of *MECOM, PRIM2*, and *MYC*, and loss of *RB1*^*7,8*^. T59 further had two pronouncedly different epithelial cell populations: (i) a larger population comprising of ≈90% of the cells, in which the majority of cells were more confidently classified as differentiated (mean margin of 0.34), and (ii) a smaller, presumably subclonal, population of cells that was characterized by additional alterations on chromosome 10, enrichment for proliferative subtype calls (p < 2.2 · 10^−16^, Fisher’s exact test), and increased subtype ambiguity (mean margin of 0.17). Tumor T90 displayed characteristics of a foldback inversion profile including a high degree of genomic rearrangements and a strong *CCNE1* amplification^7,8^; Tumor T77 had a heterogeneous copy number profile indicating presence of several subclonal cell populations. Subtype calls for the epithelial cells of both tumors (T77 and T90) were highly ambiguous between differentiated and proliferative (mean margin of 0.16 and 0.22, respectively).

**Figure 5:**
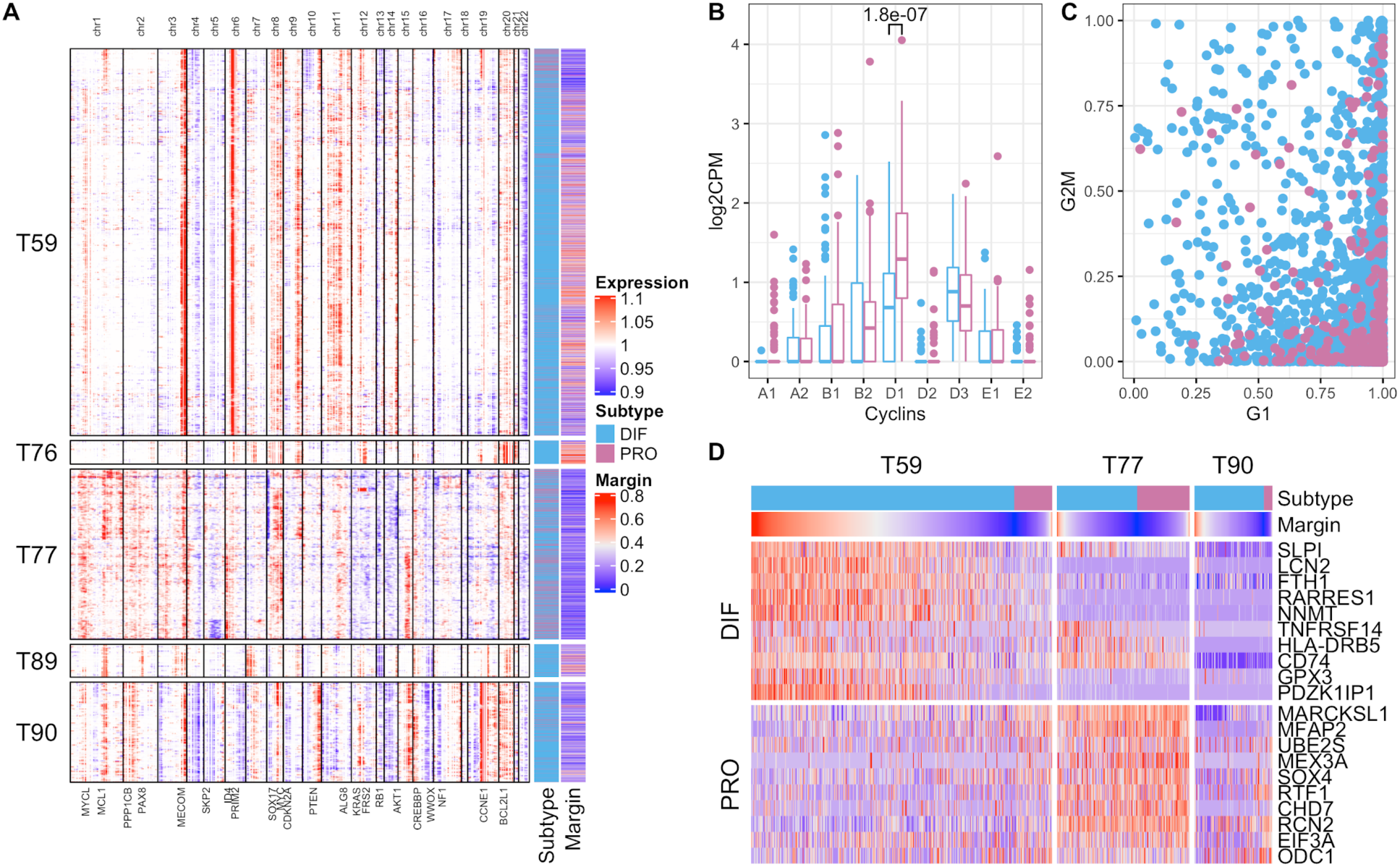
Copy number profiles, cell cycle phase, and marker gene expression of tumor epithelial cells. **(A)** Chromosomal landscape of copy number amplifications (red) and deletions (blue), inferred from scRNA-seq data using inferCNV^44^, for 6,482 individual epithelial cells of 5 tumors. Selected driver genes located in regions of recurrent bulk SCNAs of TCGA HGS ovarian tumors (Figure 2) are shown at the bottom. The annotation bars on the right show consensus subtype and classification margin (higher margin values correspond to more confident subtype assignment). **(B)** Comparison of cyclin expression in differentiated and proliferative epithelial cells of Tumor 59, displaying a significant up-regulation of Cyclin D1 in proliferative epithelial cells (*p* = 1.8 · 10^−07^, Wilcoxon rank-sum test). Shown is the comparison of the 100 cells most confidently classified as DIF and PRO, respectively, based on margin scores. **(C)** Distribution of G1 phase (*x*-axis) and G2/M phase (*y*-axis) cell cycle scores for differentiated and proliferative epithelial cells of Tumor 59 as obtained using the cyclone classifier^45^. **(D)** Cross-tumor patterns of gradual down-regulation of differentiation markers (DIF panel) and gradual up-regulation of proliferation markers (PRO panel) in tumor epithelial cells when sorting cells by subtype assignment and classification margin (annotation bars at the top).

For the three tumors with a larger number of epithelial cells captured and a substantial proportion of cells assigned to the proliferative subtype (T59, T77, and T90), we further analyzed whether differences in subtype assignment can be explained by differences in cell cycle activity. Therefore, we (i) inspected the expression levels of cell cycle phase-specific cyclins, and (ii) computationally assigned cells to different cell cycle phases using the cyclone classifier^45^. However, proliferative subtype calls could not be attributed to increased cell cycle activity as cyclin expression was largely similar between differentiated and proliferative cells (Figure 5B and Supplementary Figure S16). Exceptions were the up-regulation of cyclin D1 in proliferative cells of Tumor T59, and of cyclin E1 in proliferative cells of Tumor T90, previously found associated with high proliferation rates in ovarian cancer cell lines^46^ and ER-positive breast cancer^47^. Furthermore, systematic assignment of cells to G1, G2/M, and S phase using the cyclone classifier^45^ displayed a distortion of cell cycle scores towards the G1 phase (Figure 5C), a known hallmark of cancer^48^, but proportions of cells assigned to different cell cycle phases were not significantly different between subtypes (*p* = 0.15, proportion test, *df* = 2, Supplementary Table S2). As differences in subtype assignment thus did not seem to primarily result from differences in cell cycle activity, we analyzed whether the expression of known HGSOC bulk tumor markers^4,11^ drives classification of epithelial cells as differentiated or proliferative. When sorting cells by subtype assignment and classification margin, we observed cross-tumor patterns of gradual down-regulation of tumor differentiation markers and gradual up-regulation of proliferation markers (Figure 5D). This included down-regulation of fallopian tube marker *SLPI*^*4*^, differentiation markers *LCN2*^*49*^ and *RARRES1*^*50*^, and iron storage gene *FTH1* whose down-regulation has been previously linked with more aggressive tumor characteristics and poor prognosis^51^. This was paralleled by up-regulation of markers of increased proliferative activity such as *MARCKSL1*^*52*^, *MFAP2*^*53*^, ubiquitin conjugating enzyme *UBE2S*^*54*^, and transcription factor *SOX4* whose up-regulation is associated with the loss of epithelial features^55^ and late-stage high-grade malignant phenotypes^56^, together indicating a transition from differentiated DIF-EPI cells to de-differentiated / proliferative PRO-EPI cells.

## Discussion

We analyzed intra-tumor heterogeneity of HGSOC subtypes and investigated whether ambiguity in subtype classification can be attributed to the polyclonal composition of HGSOC tumors. This analysis addresses unresolved questions of whether proposed subtypes are early carcinogenic events that are reflected in later genomic development, whether they occur late in tumor development giving rise to subclonal expansion, or whether the notion of discrete HGSOC subtypes should be dismissed altogether. This distinction is clinically relevant because subtype-like changes occurring later in tumor development are likely to be heterogeneous in polyclonal tumors and are, thus, difficult targets for subtype-specific therapy, and furthermore, misplaced use of subtype terminology may interfere with a fuller understanding of HGSOC carcinogenesis. To infer the evolutionary timing of the development of subtype-like properties, we (i) analyzed recurrent SCNAs in TCGA tumors for subtype association, and (ii) tested whether subtype-associated SCNAs tend to predominantly occur clonally (early) or subclonally (late). We first discuss genomic characteristics of the proposed subtypes based on SCNA subtype association and subclonality, and contrast these findings with the results from the single-cell analysis of six HGSOC tumors. We then propose a more realistic model of continuous tumor development that includes mixtures of subclones and evolution between properties previously associated with discrete subtypes and discuss possible implications for chemotherapy.

### Inference of subtype evolution from somatic DNA alterations

Reliable inference of subtype evolution from bulk tumor genomics is possible because recurrent CN alterations in HGSOC are associated with the proposed transcriptome subtypes. Absolute CN analysis from SNP arrays and whole-exome sequencing allows detection of subclonal alterations^30,39^, and thus, inference of the subclonality of proposed subtypes. As reported elsewhere^39^, we also found individual CN calls and per-tumor estimates of purity and ploidy to be consistent across experimental platforms (SNP arrays and whole-exome sequencing) and computational methods (ABSOLUTE^30^ and PureCN^40^). To demonstrate this novel approach, we showed that soft tissue sarcoma (STS) subtypes are driven by clonal SCNAs that occur early in tumor development. HGSOC and STS are both characterized by recurrent SCNAs and low levels of somatic mutations^41^; comparable analysis of other cancer types that are mainly driven by somatic mutations would require incorporating allele-specific analysis of somatic mutations^30,40^. We note that this method could be extended to allele-specific analysis of point mutations for other cancer types, as long as those mutations are not too ubiquitous or too rare.

In HGSOC, proliferative tumors are associated with a large number of amplifications, higher ploidy and increased frequency of genome duplication. Amplifications associated with proliferative tumors drive a pattern that subtype-associated SCNAs tend to be more frequently subclonal than other alterations. By contrast, differentiated tumors display lower alteration frequency, close-to-normal ploidy, and smaller fractions of subclonal alterations. Thus, differentiated and proliferative tumors appear to represent different ends of an evolutionary time scale. Since tumors generally evolve from a founder clone with fewer genomic lesions to multiple clones with accumulated lesions^57,58^, these observations indicate that the proposed subtypes are more likely stages along such a process of tumor evolution.

### Single-cell sequencing reveals cell type-driven subtype assignments and presents new challenges for subtyping single cells

Single-cell analysis further suggests transitions from the differentiated towards the proliferative spectrum. Tumor epithelial cells with fewer copy number alterations are more confidently classified as differentiated, expressing signals associated with fallopian tube tissue identity and differentiation of the founder clone. Higher levels of genomic rearrangements, typically detectable in subclones, are more frequently found in epithelial cells classified as proliferative, expressing a heterogeneous set of markers associated with more aggressive tumor characteristics, loss of epithelial cell features, and poor prognosis. Single-cell analysis further confirms large differences in proportion of epithelial, immune, and stromal cells between tumors. Assignments to the immunoreactive and mesenchymal spectrum were driven by large fractions of tumor-infiltrating immune and stromal cells, respectively, consistent with lower purity and with reports linking the immunoreactive and mesenchymal subtypes to the tumor microenvironment^16,38,59^. The observation that discrete subtype calls were frequently less confident on single cells than on the bulk tumor further supports the notion that the proposed bulk subtypes are not well defined. The observed ambiguity in classification of single cells did not result from the low coverage of scRNA-seq data^43^, as downsampling of bulk RNA-seq data to match the coverage of the scRNA-seq data still allowed for a more confident subtype assignment. Instead, the bulk tumor classifier can be understood as marking (i) cell type proportions, and (ii) accumulation of mutations in cancer epithelial cells, with only the latter being relevant to classification of single epithelial cells. Ambiguity in classification of single epithelial cells reflects that the bulk tumor classifier contains many signature genes not expressed in individual epithelial cells or expressed along a continuous spectrum, as opposed to signature genes not being sufficiently expressed for detection at single-cell level. This is consistent with previous observations^11,16^ that each HGS ovarian tumor is represented by multiple expression signatures at different levels of activation which are effectively deconvolved at single-cell level. Therefore, these observations also question the utility of the proposed transcriptome subtypes for describing distinct molecular features of individual HGS ovarian cancer cells. As molecular subtypes at bulk resolution are largely determined by tumor composition with the most abundant cell type driving classification, their application does not seem to be meaningful at single-cell resolution. Although the proposed subtypes might still provide for a convenient summary of a tumor with some predictive value, our findings prompt development of a separate subtyping scheme for individual HGS ovarian cancer cells as a foundation for establishing whether true intrinsic HGSOC subtypes, characterized by different molecular features as, for instance, described for soft tissue sarcoma, exist.

### A model of HGSOC heterogeneity based on tumor evolution and composition

Rather than supporting the existence of discrete subtypes, our observations are more consistent with a model that places the differentiated and the proliferative subtype at opposite ends of the timeline of tumor development, with characteristics of differentiated tumors occurring earlier in malignancy, and characteristics of proliferative tumors occurring at a later time. Increasing genomic instability and subclonal expansion along this timeline, likely spanning several years, is consistent with previous reports^24,60^. Furthermore, (i) mean patient age at diagnosis was lowest for differentiated tumors (55 years), and highest for proliferative tumors (64 years)^13^; (ii) differentiated tumors displayed the highest levels of lymphocyte infiltration (>40%)^13^, indicating an active immune response at an earlier stage of tumor development. In contrast, proliferative tumors displayed negligible lymphocyte infiltration (<5%)^13^, consistent with an adapted tumor successfully evading the immune response at a later point in evolution^61^. Immunoreactive tumors are associated with infiltrating macrophages and different histopathological classification^62^; and mesenchymal tumors with stromal expression^16,59^ and potentially with different tissue of origin^14^. Although immunoreactive and mesenchymal tumors may thus represent different evolutionary starting points, characteristics of both are more likely to result from different cellular mixtures occuring along the same evolutionary timeline (Supplementary Discussion S3.1).

We therefore propose dismissing the model of discrete subtypes for HGSOC. While tumors of different patients display distinctive properties, we propose understanding these properties as existing on a *spectrum*, and as having the potential to change over the course of tumor development. This interpretation is consistent with the reported continuum of HGSOC genomes shaped by individual CN signatures^8^ (Supplementary Discussion S3.2), and consolidates and extends previous reports on extensive temporal and spatial heterogeneity in HGSOC^24,60^. The presence of different sources of heterogeneity presents a challenge for effective therapy, since tumor evolution over the course of therapy and relapse has been repeatedly attributed to drug-resistant subclones, which expand under the selective pressure of therapeutic intervention^24,60^ (Supplementary Discussion S3.3). Therapies targeting genomic mutations should thus focus on events occurring earlier in tumor evolution, because those occurring later will be subclonal, even if they grow to dominate the tumor. Among HGSOC SCNAs, such early clonal alterations tend to be deletions, including frequent loss of RB1, NF1 and PTEN (Supplementary Figures S4 and S6), which are, however, difficult therapeutic targets^63^.

## Conclusions

In this study, we investigated whether ambiguity of gene expression-based HGSOC subtypes results from intrinsic ambiguity at the level of individual tumor cells or from a mixture of subclones of different defined subtypes. Although subtype-associated DNA alterations tend to occur more frequently subclonal, we found this to be merely attributable to the overall higher ploidy and subclonality of tumors of the proliferative spectrum, rather than providing evidence for the existence of subclones with different well-defined subtypes. On the other hand, subtype classification on single cells demonstrated that ambiguity results from a combination of (i) mixture of cell types at bulk level, and (ii) apparently evolving characteristics from one subtype spectrum to another. From these observations we conclude that the notion of discrete subtypes does not realistically represent the heterogeneity and genomic complexity of HGSOC. We instead propose that HGSOC is characterized by heterogeneity defined primarily by tumor evolution and composition. This perspective is in agreement with recent genomic classifications of HGSOC^7,8^, reported temporal and spatial heterogeneity^24,60^, and reconciles findings from bulk tumor and single cell analysis. In this model, tumors evolve from a largely intact genome (early differentiated spectrum) towards a comprehensive loss of genome integrity (late proliferative spectrum), driven by stochastic and individually different genomic alterations from a constrained set of evolutionary moves that give rise to increasing genomic instability and subclonal expansion. Together with heterogeneity in tumor purity and composition, driving assignment of tumors to the immunoreactive and the mesenchymal spectrum, this provides an explanation for ambiguity in subtype classification that exists also on the cellular level to an extent even exceeding classification ambiguity of the bulk tumor. Experimental validation of this model using human samples will be challenging, as it is not feasible to collect longitudinal samples from a patient as their tumor evolves; longitudinal samples could be obtained from transgenic mice that develop autochthonous ovarian cancer. Molecular analysis of early-stage HGS ovarian tumors from multiple independent cohorts provides supporting evidence as the majority of these tumors have a differentiated phenotype with low numbers of SCNAs (Supplementary Table S3). As a strong relationship exists between patient age and tumor stage of HGSOC^4,64^, this also implies that an earlier stage is likely at least partly due to earlier detection, as opposed to just a more indolent tumor, and that stages likely develop over decades. With the availability of more single-cell data for HGSOC in the near future, it will also be possible to more comprehensively study tumors at critical transition stages before, during, and after treatment.

## Acknowledgments

LG was supported by a research fellowship from the German Research Foundation (GE3023/1-1 and GE3023/1-2). RLR, CH, ACN, BW, and TS were supported by grants from the Jan Chorzempa Cancer Research Endowed Fund (TKS), the Masonic Cancer Center’s Translational Working Group Grant (TKS), University of MN Grand Challenges Grant (BW, ACN, TKS), the American Cancer Society CSDG (ACN, #132574-CSDG-18-139-01-CSM) and institutional grants to the University of Minnesota from NIH/NCI P30CA07759821 and CTSI NCATS UL1TR00249402. GP was supported by grant 5P30CA006516-53, and LW by grants U24CA180996 and 1R03CA191447-01A1 from the National Cancer Institute of the National Institutes of Health.

