## Supplemental Materials for "Multi-omic analysis of subtype evolution and heterogeneity in high-grade serous ovarian carcinoma"

**Supplementary Material**

### S1 Supplementary Methods

#### S1.1 Ethical statement

All single-cell patient samples were collected using protocols approved by the University of Minnesota's Institutional Review Board (Protocol numbers 1408M52905 and 1611M99903). Controlled-access TCGA data (DNA sequencing BAM files) were downloaded from the Genomic Data Commons (GDC) using a key obtained through dbGaP (dataset phs000178.v10.p8.c1, Data Access Request #47794-4), and handled in accordance with the General Research Use requirements for use of this dataset.

#### S1.2 Single-cell patient samples

All patient samples were collected using protocols approved by the University of Minnesota's Institutional Review Board (Protocol numbers 1408M52905 and 1611M99903). Women were consented to the study if they had clinical, laboratory and/or imaging findings suspicious for ovarian cancer, including recurrent ovarian cancer. Women were enrolled upon pathological confirmation of ovarian cancer. Tumor specimens taken from the operating room were used for clinical diagnoses and excess tissue was released to the study. Samples were kept in media at 4°C. Tissue samples were dissected with scalpel and scissors to remove necrotic and fatty tissues and to dice the tumor specimens into small pieces. Tumor pieces were digested at 37°C and gentle shaking following the Miltenyi GentleMacs Dissociation system protocol (Miltenyibiotec, San Diego CA). This protocol uses a proprietary mix of enzymes and specially designed centrifuge tubes to generate a single cell suspension. Cells were filtered through a 70 micron filter, washed 2× in PBS and incubated with red blood cell lysis to remove red blood cells. Cell viability was determined based on trypan blue exclusion. Cells were transported to the University of Minnesota core sequencing facility for processing within 3 hours after excision from the patient.

#### S1.3 Single-cell RNA quantification

RNA was converted to cDNA and tagged with barcodes following the 10x Genomics Single Cell 3' Protocol utilizing the ChromiumTM Single Cell 3' v2 Library & Gel Bead Kit and ChromiumTM Single Cell A Chip Kit (10x Genomics, Pleasanton, CA). The cDNA was sequenced using an Illumina HiSeq 2500 flow cell with Rapid chemistry and 100 bp paired end sequencing, producing an average of 235 million reads per patient sample with an average of 11,502 cells/sample and 23,000 reads/cell. Fastq files were analyzed using `CellRanger` (v3.0.2) software which performs indexing, demultiplexing, and alignment to produce a matrix of RNA abundance (cells × genes) using the Unique Molecular Identifier (UMI) counts. Raw and processed scRNA-seq data is available upon request and will be deposited in GEO upon acceptance.

#### S1.4 Bulk RNA quantification

RNA was extracted from fresh-frozen tissue using the PureLink$^{TM}$ RNA Mini Kit from Invitrogen (Cat. No.12183020) following manufacturer's protocol. RNA-seq libraries were prepared using the Illumina TruSeq Stranded mRNA kit (Cat No. 20020594) following manufacturer's protocol. Libraries were sequenced on an Illumina NovaSeq instrument in a single batch with 51 bp paired end reads resulting in ≈25-30M reads per sample. FASTQ files were processed through the `PURR` RNA-seq pipeline developed by the University of Minnesota Informatics Institute/Research Informatics Services (`https://github.com/msi-ris/CHURP/wiki/PURR-Example`). Briefly, raw reads were summarized with `FastQC` [1], trimmed with `Trimmomatic` [2], and aligned to the reference genome with `HISAT2` [3]. Cleaned alignments were counted with `featureCounts` [4] in a strand specific manner. Final sample data across the cohort had a minimum of 24.4M trimmed reads, a minimum of 80% uniquely mapped reads, and a maximum of 3.6% rRNA contamination. All samples achieved a minimum of 25,000 expressed features.

#### S1.5 Single-cell data processing

Data preprocessing, quality control (QC), normalization, dimensionality reduction, and clustering was carried out based on the Bioconductor workflows for single-cell analysis [5], using functionality implemented in the `scater` [6] and `scran` [7] Bioconductor packages. Cells representing outliers for one of three QC metrics were excluded based on the median absolute deviation (MAD) from the median value of each metric across all cells. We therefore excluded cells that displayed a high proportion of reads mapped to mitochondrial genes (3 MADs), small library sizes (1 MAD), and cells with too few expressed genes (1 MAD). For the remaining cells, we excluded genes with an average expression below 0.001, i.e. with less than 1 read per 1000 cells. Normalization for differences in library size between cells was carried out using the `normalize` function from the `scater` package. Dimensionality reduction was carried out using the `denoisePCA` function from the `scran` package, which carries out principal components analysis (PCA) and removes principal components corresponding to technical noise. Clustering was carried out using the `buildSNNGraph` function from the `scran` package with the number of nearest neighbors (argument `k`) set to 25. The resulting $k$-nearest-neighbors graph was clustered using the function `cluster_walktrap` from the `igraph` package [8]. Summary statistics of the various processing steps are shown in Supplementary Table S1.

#### S1.6 Single-cell cell type annotation

Cell types were annotated based on marker gene expression and similarity to reference transcriptomes of pure cell types. Marker genes for epithelial, endothelial, lymphoctye, myeloid, and stromal cells were selected based on the work of Shih *et al.* [9]. Annotation transfer based on transcriptome similarity to pure cell types were carried out using the `SingleR` Bioconductor package [10] with two different reference databases: (i) The Human Cell Atlas [11], and (ii) a compilation of cell type transcriptomes from ENCODE [12] and BLUEPRINT [13].

#### S1.7 Single-cell CN profiles, cell cycle activity, and marker gene expression

Copy number profiles were inferred from scRNA-seq data using `inferCNV` [14, 15]. Input files were prepared according to the instructions under `https://github.com/broadinstitute/inferCNV/wiki/File-Definitions`, providing (i) a matrix (genes × cells) storing the raw read counts, (ii) annotations for the cells grouping cells into observations (tumor epithelial cells) and normal reference cells (myeloid, lymphocytes, stromal, and endothelial cells), and (iii) a gene order file providing the chromosomal location of each gene according to the UCSC hg38 genome assembly. Differences in cell cycle activity were analyzed by comparing the expression levels of cell cycle phase-specific cyclins in tumor epithelial cells classified as differentiated or proliferative. Comparison of cyclin expression between differentiated and proliferative cells was carried out by (i) taking all epithelial cells of a tumor into consideration, and (ii) for a restricted set of epithelial cells that were most confidently classified as either differentiated or proliferative. Significance assessment of differences in mean cyclin expression levels between subtype groups was carried out using Wilcoxon's rank-sum test. Systematic assignment of cells to different cell cycle phases was carried out using the `cyclone` classifier [16] as implemented in the function `cyclone` from the `scran` package [7].

#### S1.8 Subtype classification on scRNA-seq data from Winterhoff et al., 2017

Bulk and single-cell RNA-seq data for one fresh HGSOC specimen was obtained from the Supplementary Material of Winterhoff *et al*, 2017 [17]. Subtypes were classified using the `consensusOV` package [18, 19]. The 800-gene signature was derived by selecting the 200 most representative genes per subtype based on differential expression [20]. TCGA bulk RNA-seq and microarray data were obtained using the `curatedTCGAData` Bioconductor package [21]. Downsampling of TCGA bulk RNA-seq data to match the coverage of the scRNA-seq data was carried out using the function `downsampleMatrix` of the `DropletUtils` package [22].

### S2 Supplementary Results

#### S2.1 Soft tissue sarcoma as a negative control

HGSOC and adult soft tissue sarcoma (STS) are both characterized by low frequency of somatic mutations, but high frequency of SCNAs [23]. In contrast to TCGA ovarian carcinomas, which are exclusively of high-grade serous type, TCGA ST sarcomas include several major types, each characterized by specific genomic features, as expected for true intrinsic subtypes. Transcriptome clustering of 259 TCGA ST sarcomas [23] was largely determined by STS type with (i) one cluster exclusively composed of Leiomyosarcoma (LMS), (ii) another cluster dominated by dedifferentiated liposarcoma (DDLPS), and (iii) the third cluster mostly consisting of undifferentiated pleomorphic sarcoma (UPS) and myxofibrosarcoma (MFS), two molecularly related STS types [24]. When testing a total of 64 SCNAs (23 amplifications / 41 deletions) for association with the transcriptome clusters, we found a strong enrichment of nominal p-values near zero, corresponding to 41 of 60 alterations being significantly associated (FDR $<0.1$, Supplementary Figure S2b). Association with the STS type-dominated transcriptome clusters was negatively correlated with subclonality prevalence (Figure 3D of the main manuscript), consistent with the assumption of these being intrinsic STS type-specific events[23, 25]. Regions of strongest subtype association and concomitantly low subclonality included the *MDM2* amplification (Supplementary Figure S4), a key driver of DDLPS and MFS/UPS, but rarely occurring in LMS. A notable exception from the observed trend was the *TP73*-containing telomeric deletion occurring predominantly subclonal in DDLPS and MFS/UPS, yet predominantly clonal in LMS (Supplementary Figure S4).

#### S2.2 Transcriptome subtyping of 66 stromal and epithelial cells

Given the highly consistent subtype calls for TCGA tumors assayed by bulk RNA-seq and microarray ( S7A), we next sought to determine the suitability of applying the `consensusOV` classifier [18] to scRNA-seq data. Therefore, we first applied the consensus classifier to 44 epithelial cells and 22 stromal cells of an HGSOC tumor for which a recent study reported heterogeneity within ovarian cancer epithelium and cancer-associated stromal cells [17]. As shown in Supplementary Figure S7C, the majority of epithelial cells were classified as immunoreactive, in agreement with the classification of the bulk tumor (Probabilities of IMR: 65%, DIF: 16%, MES: 14%, PRO: 5%). However, it should be noted that the subtype calls for individual cells are not directly comparable with the subtype call for the bulk tumor, given that immune cells present in the bulk tumor were removed from the analysis of individual cells by Winterhoff *et al.* [17]. Several cells assigned to the stromal group by Winterhoff *et al.* [17] were classified as mesenchymal, consistent with the low purity of mesenchymal tumors (Supplementary Figure S1a) and observations that stromal expression drives classification of mesenchymal tumors [26].
Classification margin scores, i.e. the difference between the top two subtype scores, were systematically lower for individual cells ($24\pm15\%$) than for the bulk tumor (48%, Supplementary Figure S7B). Inspecting subtype classification probabilities of epithelial cells classified as immunoreactive ( *ClassProb* bars in Supplementary Figure S7C) revealed the differentiated subtype to often closely placing second; and vice versa for the four epithelial cells classified as differentiated. To analyze whether the observed ambiguity in classification of single cells can be explained by zero-inflation of scRNA-seq data, rendering parts of the 100-gene signature of the consensus classifier not informative, we used an extended signature of 800 genes. However, similar subtype calls (Supplementary Figure S8) and margin score distribution (Supplementary Figure S7B) indicated that the 100-gene signature already captures subtype-specific expression of single

cells. In a complementary analysis, we downsampled the TCGA bulk RNA-seq data to match the coverage of the scRNA-seq data. Classification margin scores on the downsampled bulk data resembled the distribution of the original data, and exceeded the margin scores of the scRNA-seq data (Supplementary Figure S7B).

Discrepancies of bulk and single cell subtypes reported here compared to those reported by Winterhoff *et al.* [17] likely result from (i) using different subtype classifiers, and (ii) subtype ambiguity of HGSOC bulk tumors and the observed even greater subtype ambiguity of single cells. Winterhoff *et al.* [17] applied an ad-hoc adaptation of the original TCGA microarray classifier [27], for which the authors reported substantial discrepancies when comparing subtype assignments for TCGA tumors assayed by both RNA-seq and microarray (30% of TCGA tumors with different subtype calls). On the other hand, we used the consensus classifier [18], implemented in the `consensusOV` Bioconductor package [19], previously trained on tumors concordantly classified by the TCGA classifier and two other major subtype classifiers across 15 microarray datasets, which also displayed high concordance when classifying TCGA tumors assayed by bulk RNA-seq and microarray (Supplementary Figure S7A).

### S3 Supplementary Discussion

#### S3.1 Patient prognosis

Given the apparent comprehensive loss of genome integrity, it seems initially counterintuitive that tumors of the proliferative spectrum have lower risk than mesenchymal tumors with respect to overall survival [18]. However, several studies have reported extreme levels of genomic instability to be associated with improved response to therapy[28, 29]. The mesenchymal spectrum might, thus, represent a specific genomic constellation at which patients are particularly at risk, whereas increased genomic instability and subclonality of proliferative tumors might have beneficial consequences for their prognosis. On the other hand, differences in prognosis might also be explained by origination from the fallopian tube (FT) versus ovarian surface epithelium (OSE) instead of, or in combination with, genomic differences [30]. A recent study reported that mesenchymal tumors display OSE-like expression patterns, whereas tumors of the other three subtypes display FT-like expression [31]. This was also apparent when analyzing consensus subtypes (Supplementary Figure S17) and is consistent with FT-marker expression for differentiated tumors [27]. OSE origin, stromal expression, and mesenchymal calls are thus associated with each other and with poor prognosis, making it difficult to distinguish whether site of origin, tumor composition, and/or tumor genomics is closer to the cause of the poor prognosis.

#### S3.2 Subtype association of CN signatures

Macintyre *et al.* [32] identified seven copy number signatures shaping a continuum of HGSOC genomes and described key associations and proposed mechanisms for these signatures.

From association analysis between the seven CN signatures and consensus subtype assignment for TCGA HGS ovarian tumors, we found tumors assigned to the proliferative spectrum overrepresented in tumors displaying high exposure levels of CN signature 6 (Supplementary Figure S19). Macintyre *et al.* [32] reported correlation with age at diagnosis for CN signature 6, in agreement with increased patient age for proliferative tumors that we reported before [18]. According to Macintyre *et al.* [32], CN signature 6 is also characterized by extensive focal amplifications, consistent with overrepresentation of associations between recurrent focal amplifications and the proliferative spectrum (Figure 2 of the main manuscript), and in general higher ploidy for tumors of the proliferative spectrum (Supplementary Figure S1).

Supplementary Figure S19 also shows depletion of signature 4 for proliferative tumors. According to Macintyre *et al.* [32], exposure levels of signature 4 are higher for tumors with mutated *MYC*, consistent with decreased alteration frequency of *MYC* in proliferative tumors that we (Supplementary Figure S4)

and others [27] observed before.
Although not observed for the subset of TCGA tumors analyzed in Supplementary Figure S19, it can further be speculated that good overall survival signatures 3 and 7, supposedly driven by BRCA1/2-related and non-BRCA1/2 related homologous recombination deficiency, might be especially relevant for tumors of the differentiated and immunoreactive spectrum, given their good prognosis, anti-correlation with age at diagnosis for signature 3, and previously reported association of BRCA1 disruptions with the immunoreactive subtype [33].

#### S3.3 Implications for chemotherapy

Previous studies modeling HGSOC tumor evolution over the course of therapy attributed tumor relapse to drug-resistant subclones that are originally present in the primary tumor, and that expand under the selective pressure of therapeutic intervention [34, 28]. However, surgery or biopsy is rarely performed for recurrent HGS ovarian tumors, restricting examination of relapsed disease to tumor cells from ascites [35]. From subtype analysis of 13 primary tumor samples and associated recurrent ascites samples from a recent HGSOC chemoresistance study [35], we found several recurrent ascites samples being assigned to a different subtype spectrum than the primary tumor (Supplementary Figure S18A). However, we also observed such differences in subtype assignment when comparing primary tumor and primary ascites samples (Supplementary Figure S18B), questioning the suitability of ascites samples for meaningful comparison of pre- and post-treatment subtype. Nevertheless, subtype changes after neoadjuvant chemotherapy were also reported by a recent pilot study [36], demonstrating that chemotherapy-induced changes in subtype assignment are in principle possible, likely as a result of changes in tumor composition. These findings further argue against the existence of intrinsic HGSOC subtypes that recur in a consistent and subtype-specific manner. Instead, it seems that chemotherapy differentially affects certain clones of heterogeneously composed HGSOC tumors, leading to changes in tumor composition and surrounding microenvironment [37, 38, 39]. The presence of, and interplay between, the different sources of heterogeneity presents a challenge for effective therapy, especially any therapy that targets individual tumor genomics. Thus, therapies targeting genomic mutations should focus on those occurring earlier in tumor evolution, because those occurring later will be subclonal, even if they grow to dominate the tumor. Among HGSOC SCNAs, early clonal alterations tend to be deletions, including frequent loss of *RB1*, *NF1* and *PTEN* by gene breakage events (Supplementary Figures S4 and S6).

**Table S1: Clinical characteristics and summary statistics for the five 10x tumors.** All samples were taken from omentum. CT: chemotherapy. The number of cells, genes, principal components, and cell clusters are shown as obtained after carrying out the processing steps described in Supplementary Methods S1.5

| | Tumor59 | Tumor76 | Tumor77 | Tumor89 | Tumor90 |
|---|---|---|---|---|---|
| Tumor stage | IV | IIIC | IVb | IV | IVa |
| Tumor grade | 3 | 3 | 3 | 3 | 3 |
| CT response | resistant | refractory | resistant | sensitive | sensitive |
| Cells | 13 040 | 13 747 | 6 903 | 4 933 | 3 630 |
| Genes | 18 079 | 17 416 | 18 543 | 17 429 | 18 011 |
| Principal components | 28 | 29 | 13 | 25 | 20 |
| Clusters | 12 | 10 | 13 | 15 | 10 |

**Table S2: Cell cycle phase assignments.** Numbers of epithelial cells assigned to the different cell cycle phases using the `cyclone` classifier [16] are shown for the three 10x tumors with a substantial proportion of cells assigned to both the differentiated and the proliferative subtype. Testing for differences in cell cycle phase assignments between subtypes was carried out using $\chi^2$ test, $df = 2$.

| | Tumor59 | | Tumor77 | | Tumor90 | |
|---|---|---|---|---|---|---|
| | DIF | PRO | DIF | PRO | DIF | PRO |
| G1 | 2810 | 415 | 2127 | 598 | 722 | 93 |
| G2/M | 96 | 7 | 48 | 9 | 46 | 3 |
| S | 110 | 13 | 37 | 5 | 26 | 3 |
| $p$-value | 0.15 | | 0.16 | | 0.51 | |

**Table S3: Consensus subtypes in three independent cohorts of HGS ovarian carcinomas stratified by tumor stage.** At publication time [27], the **TGCA** HGSOC cohort comprised 489 clinically annotated stage II-IV tumor samples; the final Firehose run comprises 561 stage I-IV samples. Stage I tumors are disproportionately often classified as DIF (10 of 16, 62.5%, $p = 0.009$, Fisher's exact test). Among early stage I-II tumors, 34 tumors are classified as either DIF or IMR (81%, $p = 0.002$). These findings for the TCGA cohort were replicated in two additional cohorts: (i) in a recent study by **Engqvist** *et al.*, 2018 [40], which also displays an enrichment of stage I tumors classified as DIF (11 / 29, 37.9%), and disproportionately many stage I-II tumors classified as DIF or IMR (34 / 50, 68%); and (ii) in the **curatedOvarianData** collection of 15 microarray datasets [41], that also showed an over-representation of stage I tumors classified as DIF (24 / 52, 46.2%, $p = 0.008$, Fisher's exact test), and disproportionately many stage I-II tumors classified as DIF or IMR (75 / 95, 78.9%, $p = 5.7 \cdot 10^{-5}$).

| | TCGA | | | | Engqvist18 | | curatedOvarianData | | | |
|---|---|---|---|---|---|---|---|---|---|---|
| | I | II | III | IV | I | II | I | II | III | IV |
| DIF | 10 | 6 | 120 | 27 | 11 | 3 | 24 | 12 | 388 | 89 |
| IMR | 3 | 15 | 121 | 22 | 8 | 12 | 19 | 20 | 423 | 74 |
| MES | 0 | 0 | 102 | 26 | 3 | 2 | 3 | 5 | 340 | 83 |
| PRO | 3 | 5 | 91 | 10 | 7 | 4 | 6 | 6 | 245 | 29 |

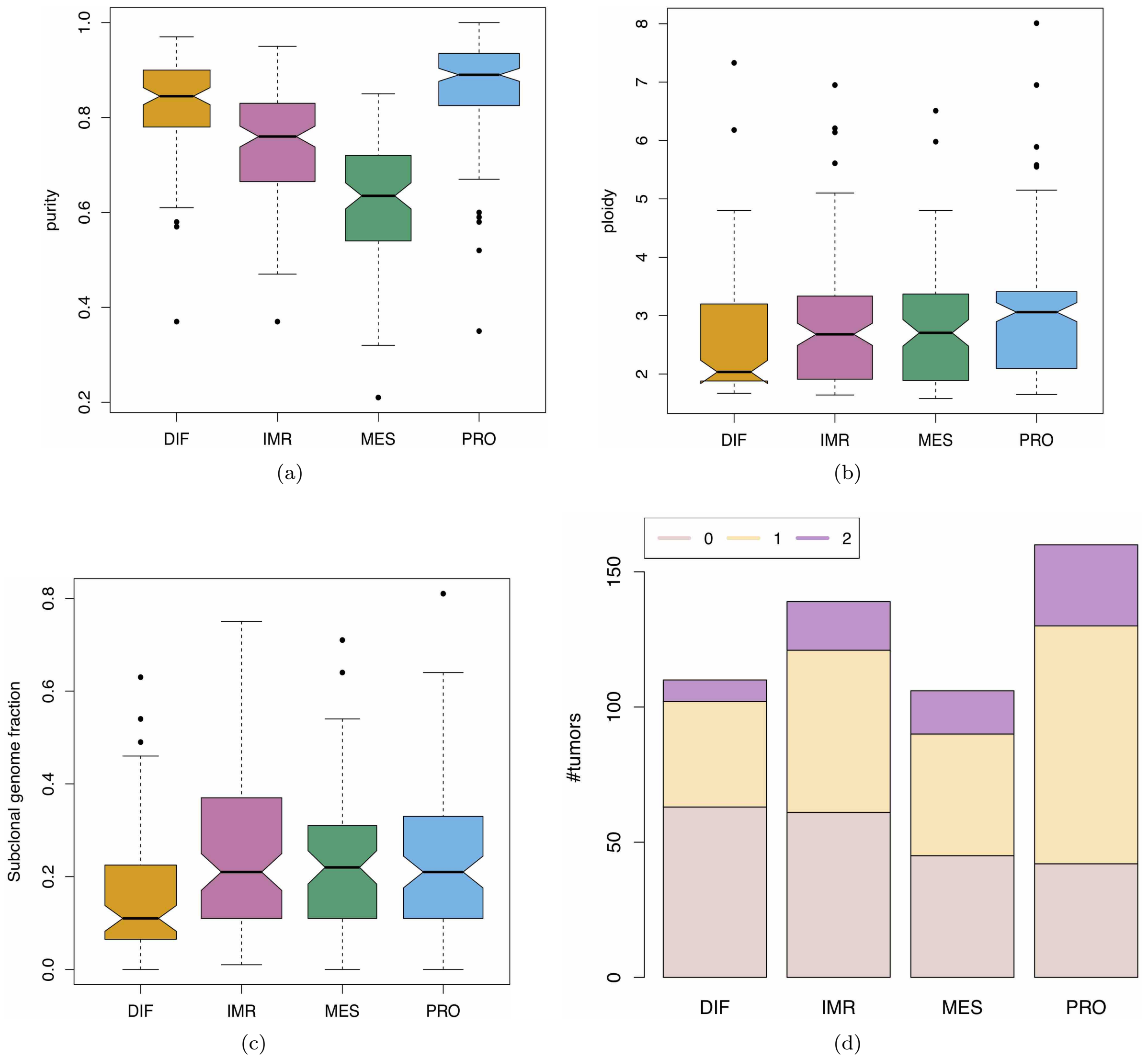

**Figure S1: Tumors of differentiated subtype are characterized by high purity, but significantly lower ploidy, subclonality, and number of genome doublings than the other three subtypes.** Subtype-stratified **(a)** purity, **(b)** ploidy, **(c)** subclonal genome fraction, and **(d)** number of genome doublings (0, 1, or 2) of TCGA HGS ovarian tumors as inferred by `ABSOLUTE`. Statistical significant differences in purity ($p < 2^{-16}$), ploidy ($p = 0.0026$), and subclonal genome fraction ($p = 0.0041$) were observed between subtypes using one-way ANOVA. The number of genome doublings also differed significantly between subtypes ($p = 8.2^{-5}$, $\chi^2$ test, $df = 6$).

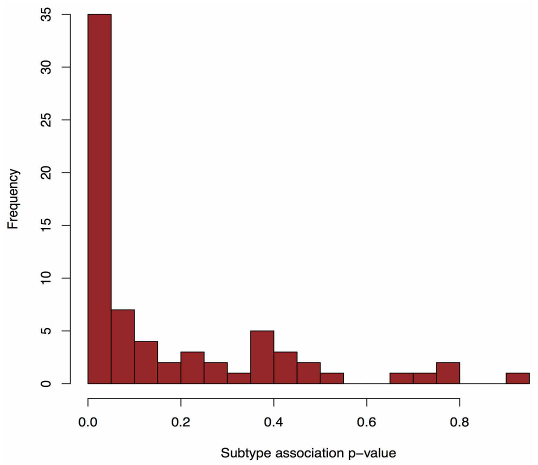

(a) OV

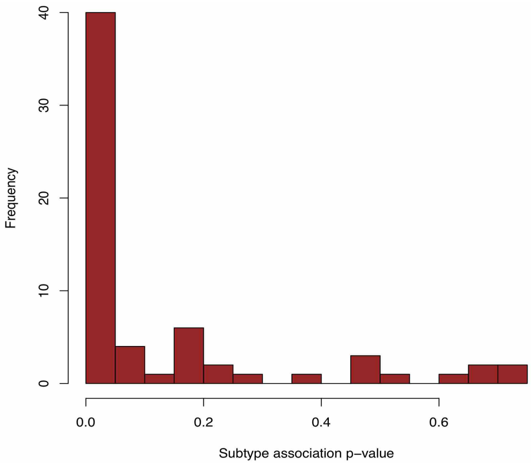

(b) SARC

**Figure S2: Association of SCNAs with transcriptome clusters.** Shown is the distribution of nominal $p$-values as resulting from testing ($\chi^2$ test with $df = 6$) recurrent focal SCNAs detected with `GISTIC2` for association with the TCGA transcriptome clusters for **(a)** 516 HGS ovarian tumors (70 SCNAs), and **(b)** 259 soft tissue sarcomas (64 SCNAs). See also Figure 3 of the main manuscript.

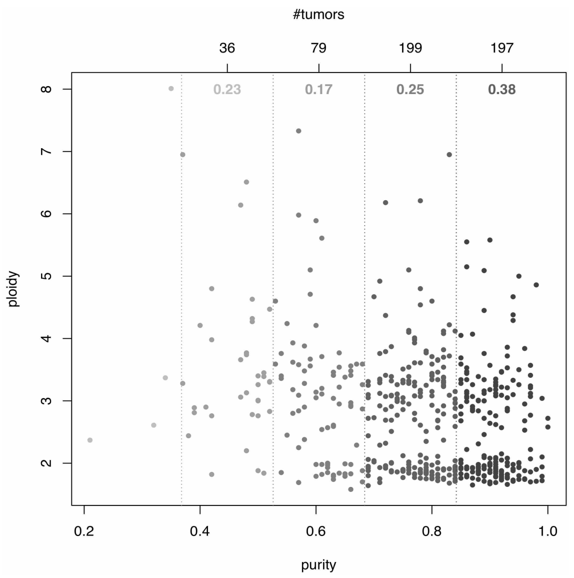

(a) equal bins

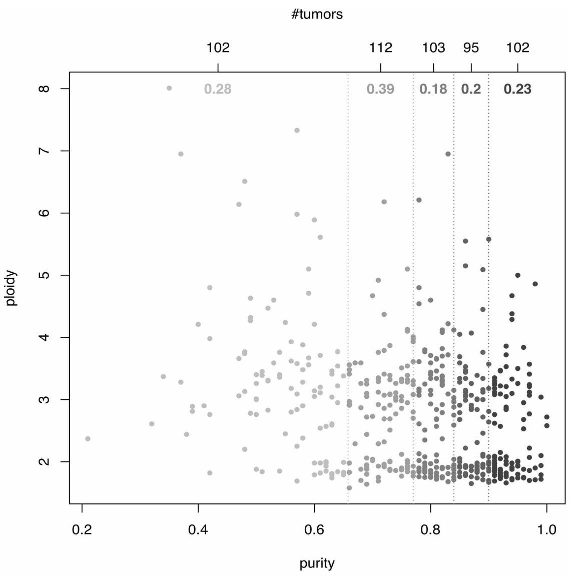

(b) quintiles

**Figure S3: Purity-stratified analysis.** The scatter plots show purity ($x$-axis) and ploidy ($y$-axis) of 516 HGS ovarian tumors as stratified by purity in **(a)** equal bins, and **(b)** quintiles. The number of tumors in each stratum is indicated on top of each plot. The correlation of subtype association and subclonality $\rho(S_A, S_C)$ is shown at the top of each stratum, except for the first stratum in **(a)** due to insufficient sample size (4 tumors).

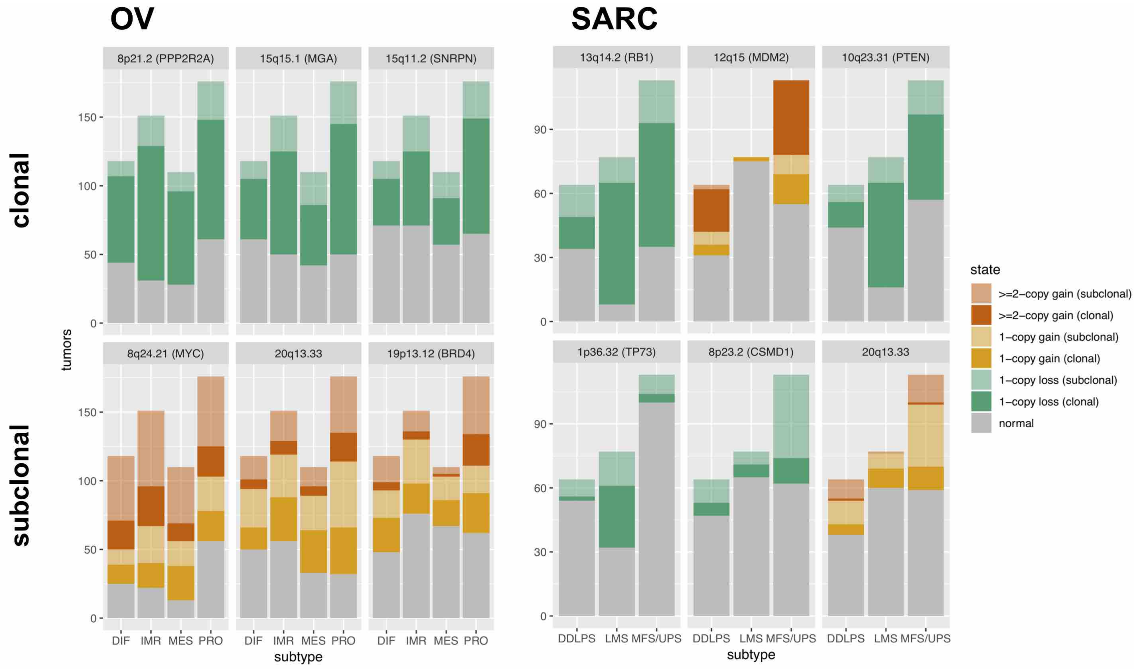

**Figure S4: Predominantly clonal or subclonal copy number alterations.** The barplots illustrate individual subtype-associated `GISTIC2` regions from Figure 3C,D of the main manuscript that occur predominantly clonal (top panel, solid color) or subclonal (bottom panel, transparent color) in TCGA HGSOC (left) or STS (right) cases. Each individual barplot displays the number of tumors ($y$-axis) of particular subtype ($x$-axis) that carry either a 1-copy loss (green), 1-copy gain (yellow), or $\geq$2-copy gain (red) in the region indicated at the top of each plot.

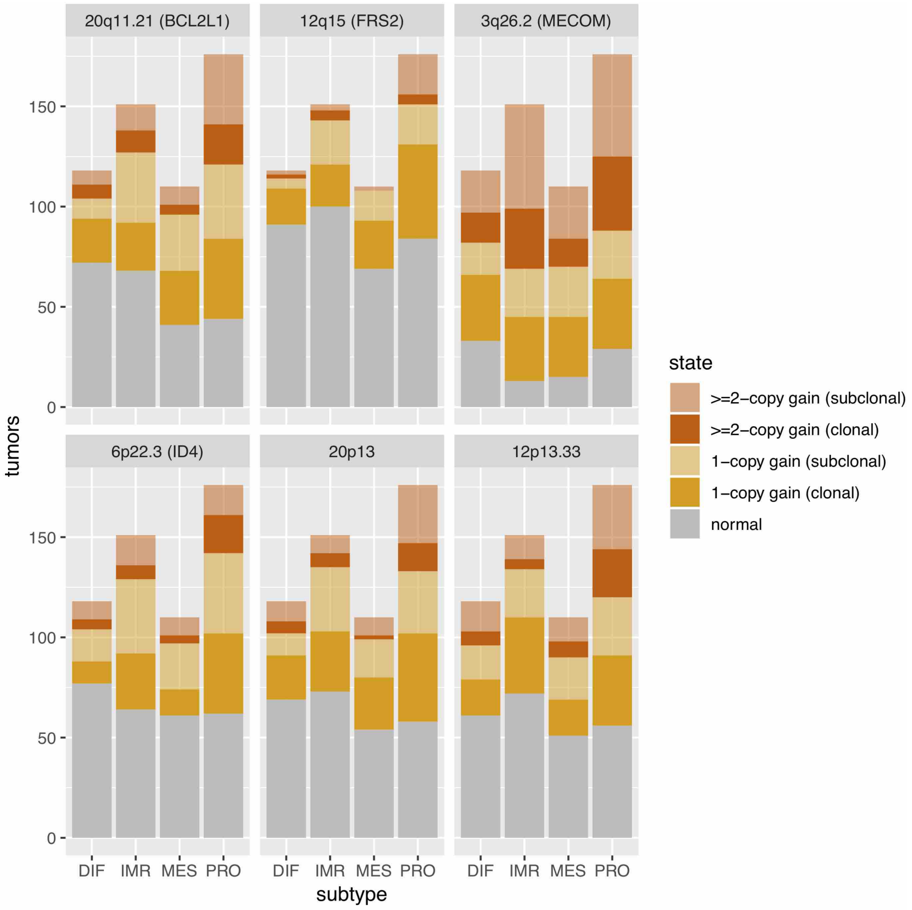

**Figure S5: Top subtype-associated regions.** The barplots illustrate individual `GISTIC2` regions of strongest subtype association from Figure 3C of the main manuscript. Each individual barplot displays the number of tumors ($y$-axis) of particular subtype ($x$-axis) that carry either a 1-copy gain (yellow) or $\geq$2-copy gain (red) in the region indicated at the top of each plot.

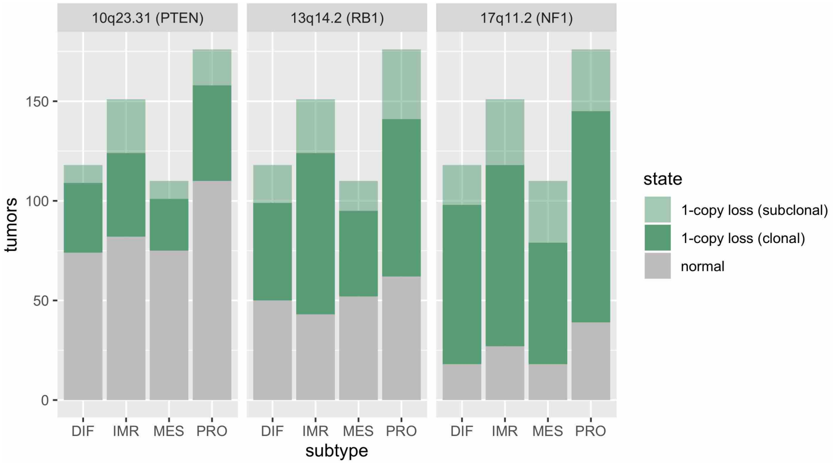

**Figure S6: Regions of frequent loss.** The barplots illustrate previously reported regions of frequent loss. As in Figure S5, each individual barplot displays the number of tumors ($y$-axis) of particular subtype ($x$-axis) that carry a 1-copy loss (green) either clonal (solid color) or subclonal (transparent color).

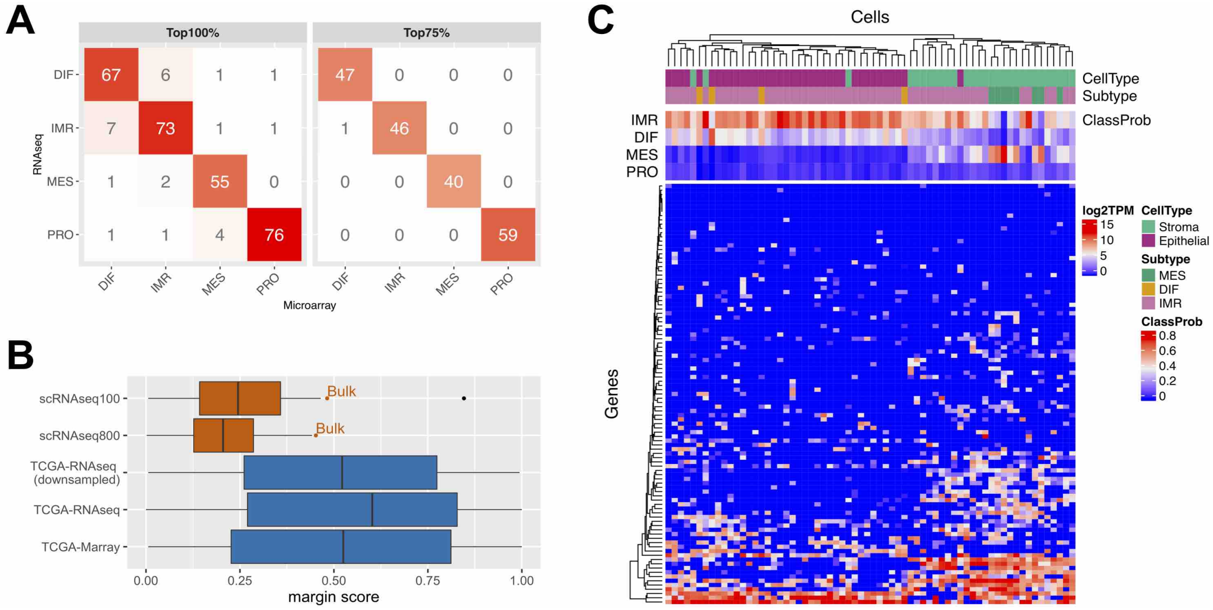

**Figure S7: Single cell subtyping.** **(A)** shows resulting subtype calls when applying the `consensusOV` [18] classifier to TCGA HGS ovarian tumors for which transcriptomic quantification was available via microarray ($x$-axis) and RNA-seq ($y$-axis). Concordance of subtype calls is shown for all tumors (top 100%, left panel) and the top 75% based on margin scores (right panel). The boxplots in **(B)** show margin score distributions of the subtype classification on 66 single cells (red) from Winterhoff *et al.* [17] as compared to the TCGA bulk classifications (blue) analyzed in (A). Single cell margins are shown when using the default 100-gene signature of the consensus classifier (*scRNAseq100*, detailed in (C)) and the extended 800-gene signature (*scRNAseq800*, detailed in Supplementary Figure S8). The margin score of the bulk tumor is indicated in both cases by the red dot. The blue box named *TCGA-RNAseq (downsampled)* shows the margin score distribution when downsampling TCGA bulk RNA-seq data to match the coverage of the scRNA-seq data. The heatmap in **(C)** depicts log2 TPM expression values of the 100-gene signature used by the consensus classifier (rows) across 66 cells (columns). The annotation bars at the top show resulting subtype calls (*Subtype*), previous characterization of cells as epithelial or stromal by Winterhoff *et al.* [17] (*CellType*), and classification probabilities for each subtype (*ClassProb*).

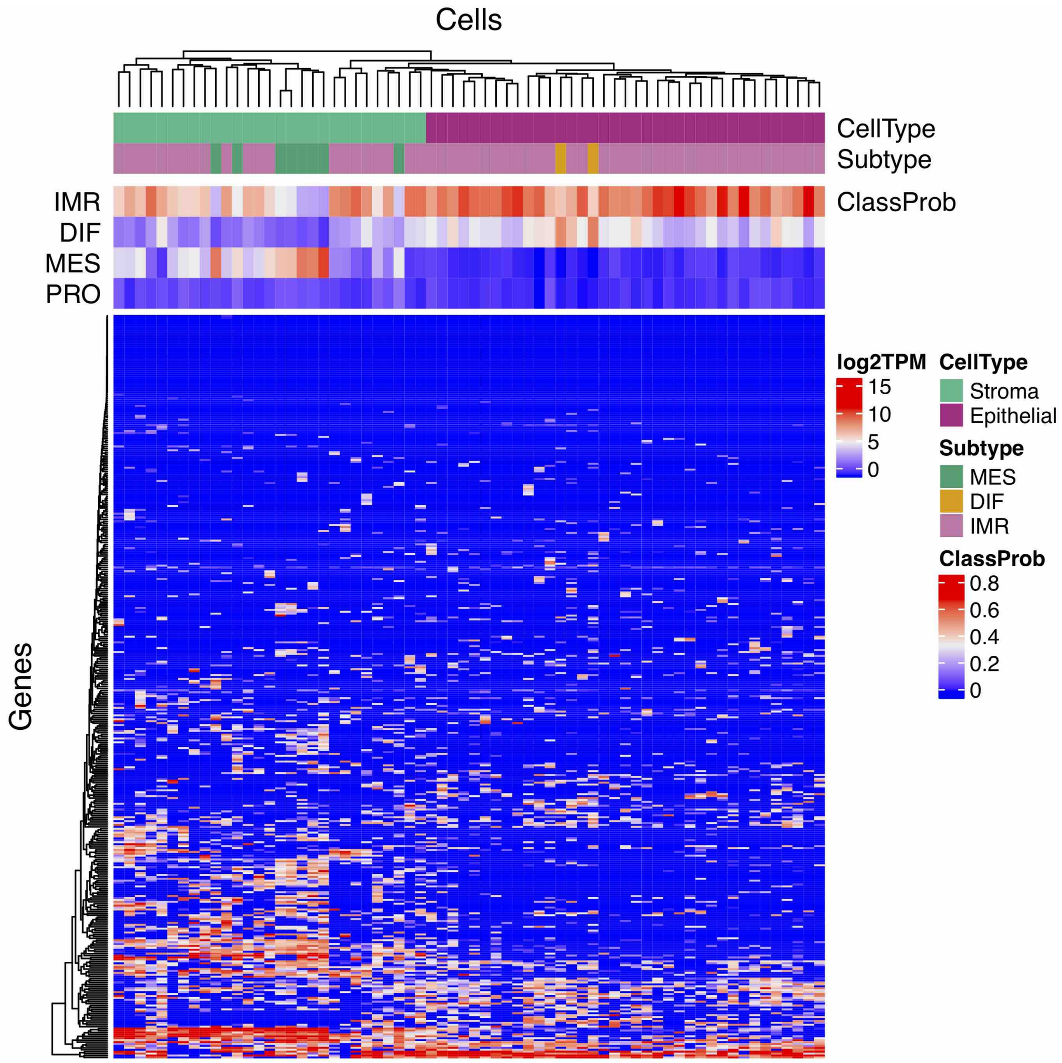

**Figure S8: Single cell subtyping with 800 genes.** The heatmap depicts log2 TPM expression values of the extended 800-gene signature (rows) across 66 cells (columns). The annotation bars at the top show resulting subtype calls (*Subtype*), previous characterization of cells as epithelial or stromal (*CellType*), and classification probabilities for each subtype (*ClassProb*).

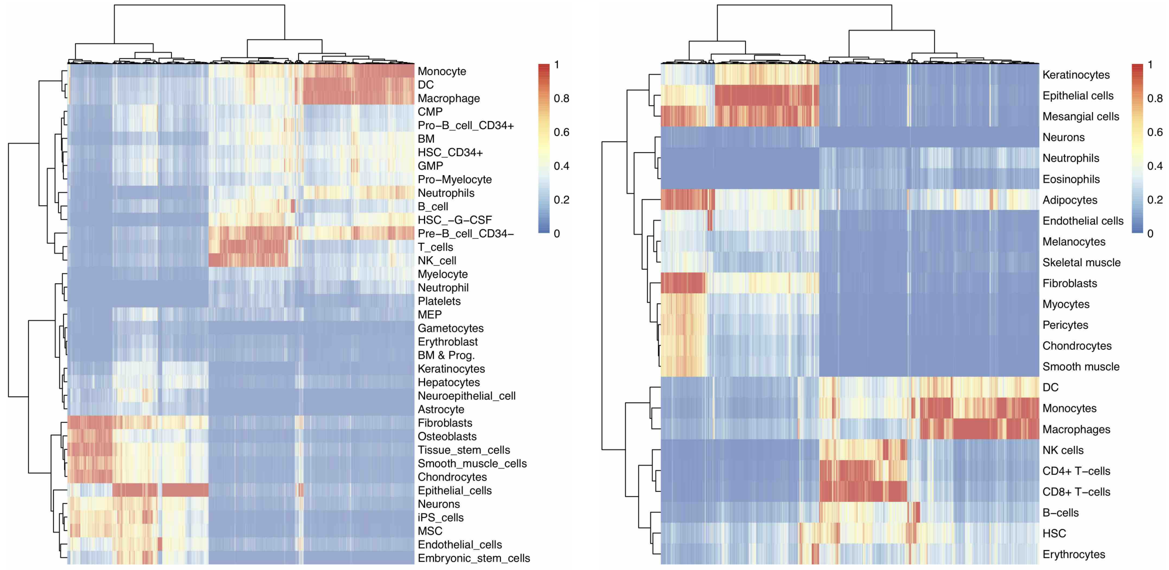

(a) Human Cell Atlas

(b) ENCODE-BLUEPRINT

**Figure S9: Cell type annotation via annotation transfer.** Cell types were annotated using `SingleR` [10] via annotation transfer from **(a)** The Human Cell Atlas [11] and **(b)** ENCODE [12] and BLUEPRINT [13]. The heatmaps show annotation scores for high-scoring cell types (rows) across 13,040 individual cells (columns) of Tumor59. Annotation transfer from both databases support distinct cell clusters of myeloid (monocytes, macrophages, dendritic cells DC), lymphocyte (natural killer NK cells, T cells, B cells), stromal (fibroblasts, pericytes, adipocytes), epithelial, and endothelial cells. Cell type assignments were also in agreement with marker gene expression (Supplementary Figures S10 and S11).

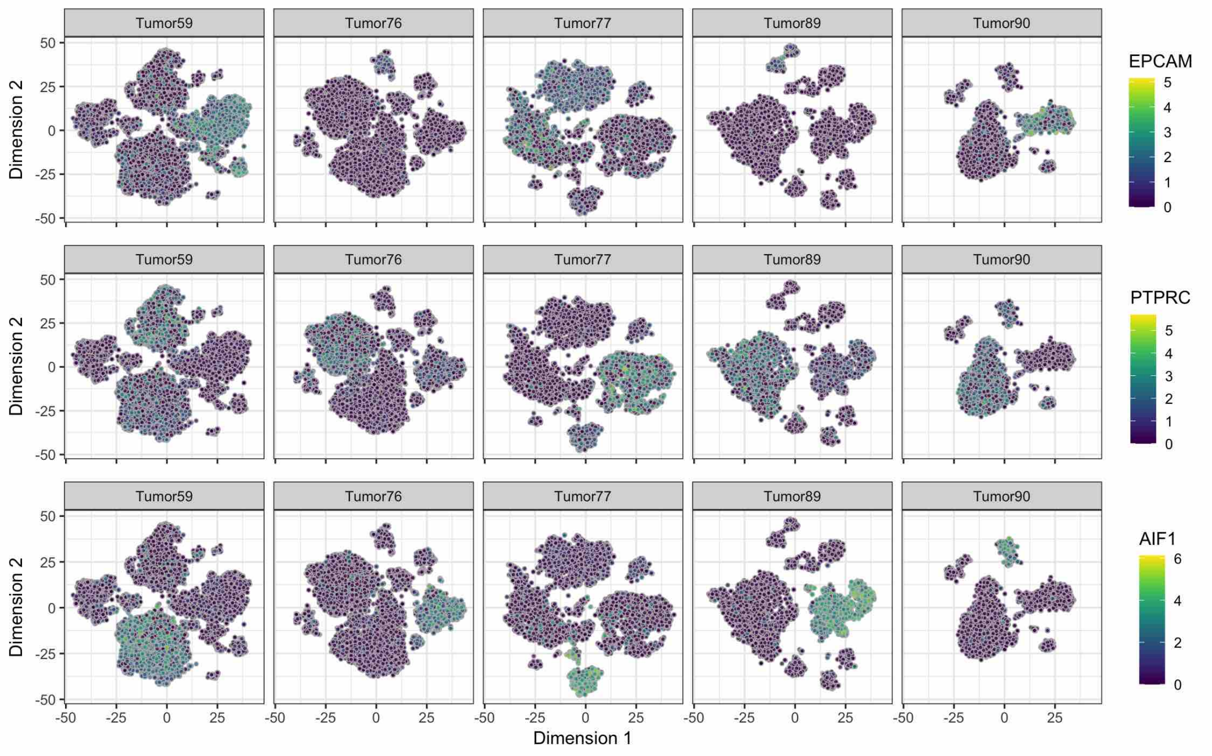

**Figure S10: Epithelial, lymphocyte, and myeloid marker gene expression.** The $t$-SNE plots show marker gene expression for epithelial ($EPCAM$), lymphocyte ($PTPRC$), and myeloid ($AIF1$) cells. Marker genes were selected based on Shih *et al.* [9]. The color scale on the right indicates the expression level given as normalized log read counts.

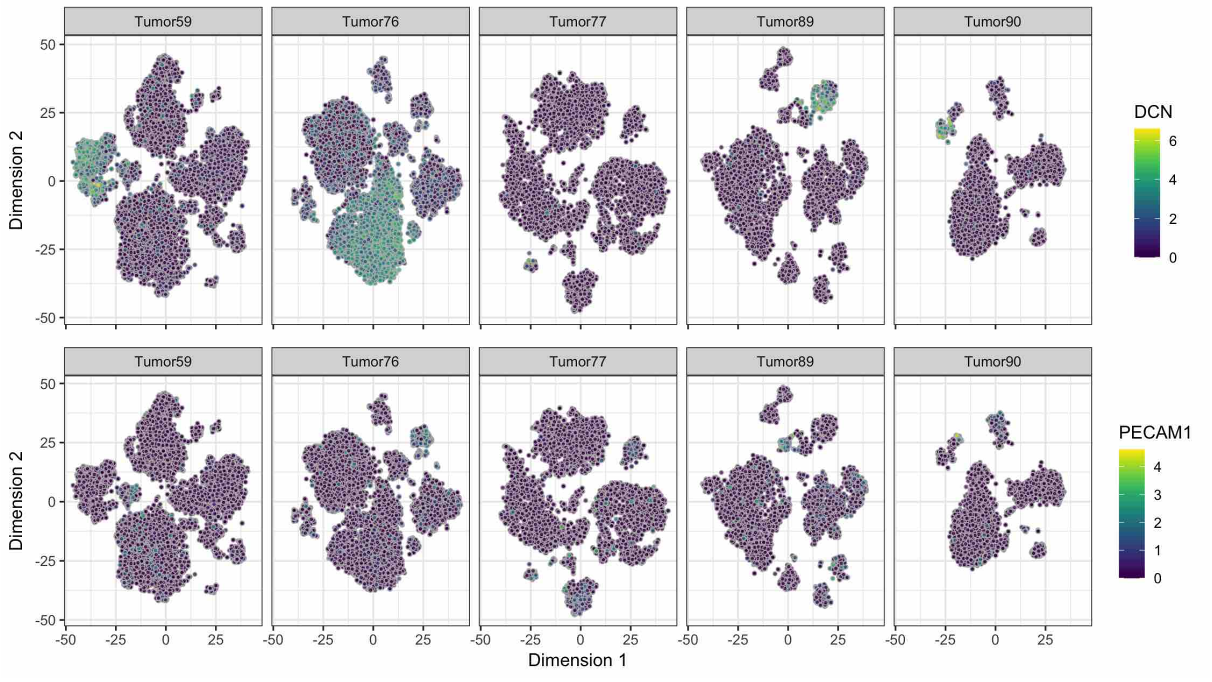

**Figure S11: Stromal and endothelial marker gene expression.** The $t$-SNE plots show marker gene expression for stromal (*DCN*) and endothelial (*PECAM1*) cells. Marker genes were selected based on Shih *et al.* [9]. The color scale on the right indicates the expression level given as normalized log read counts.

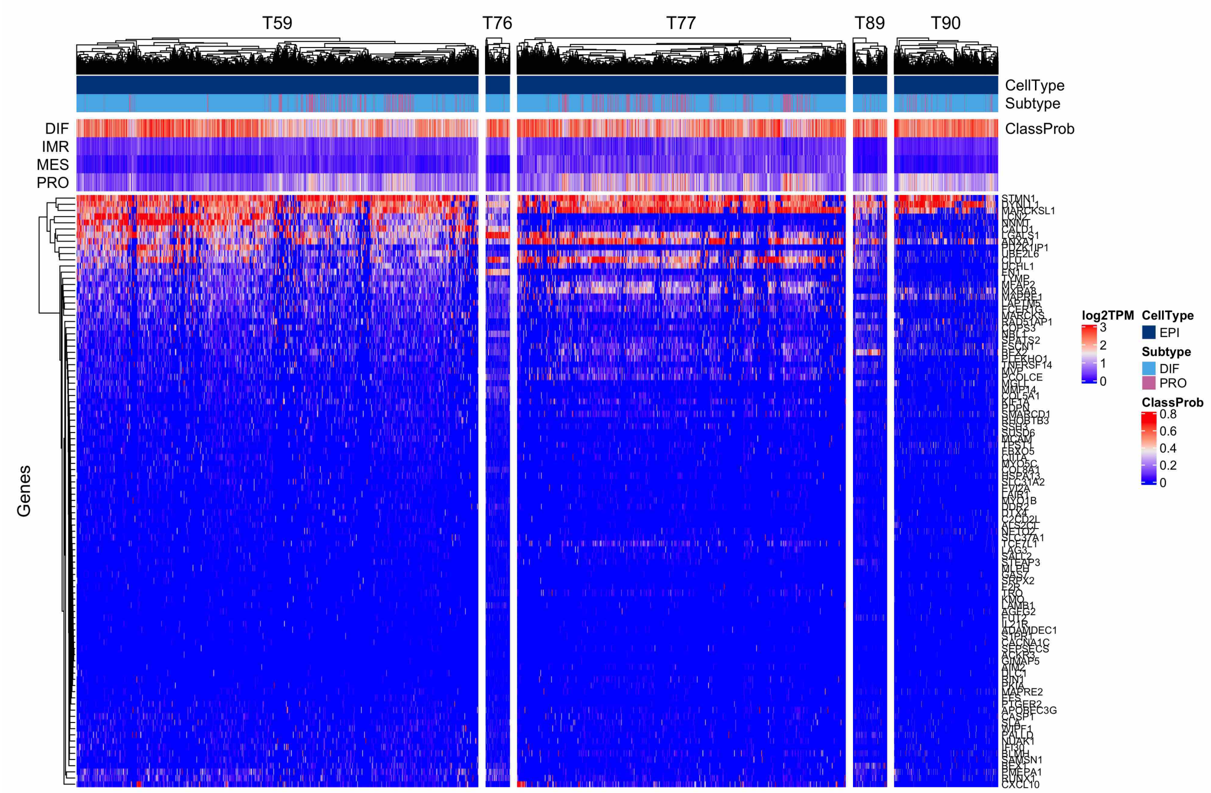

**Figure S12: Transitional patterns between DIF and PRO subtype spectrum.** The heatmaps show log2 TPM expression values of the 100-gene signature used by the consensus classifier (rows) across 3,451 / 210 / 2,824 / 293 / 893 epithelial cells of tumors T59 / T76 / T77 / T89 / T90 (columns). The annotation bars at the top show resulting subtype calls (*Subtype*) and classification probabilities for each subtype (*ClassProb*).

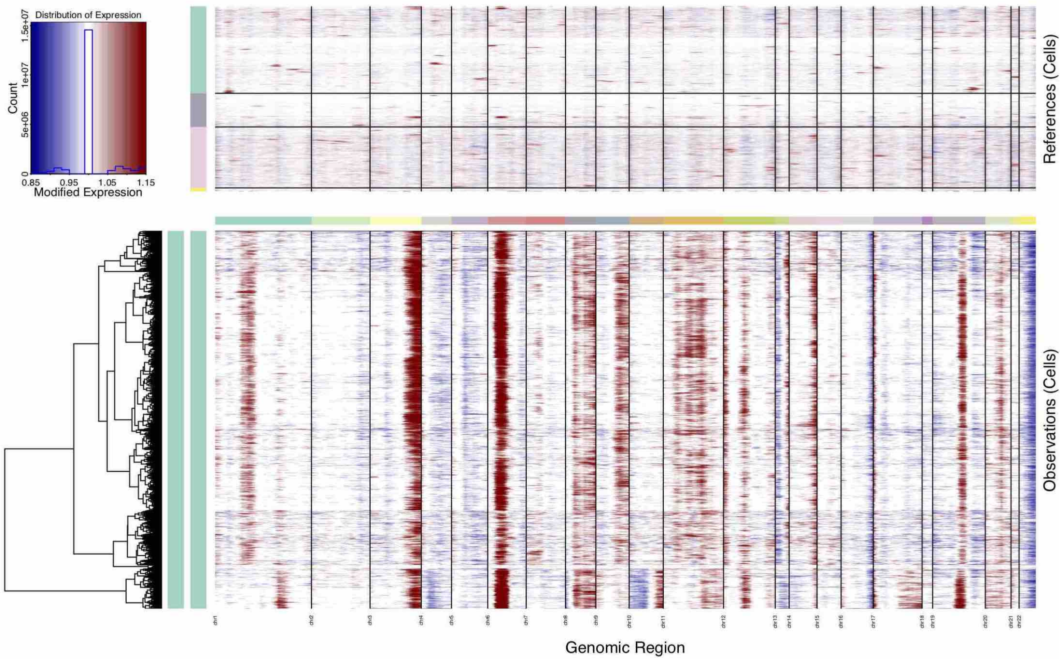

(a) Tumor T59

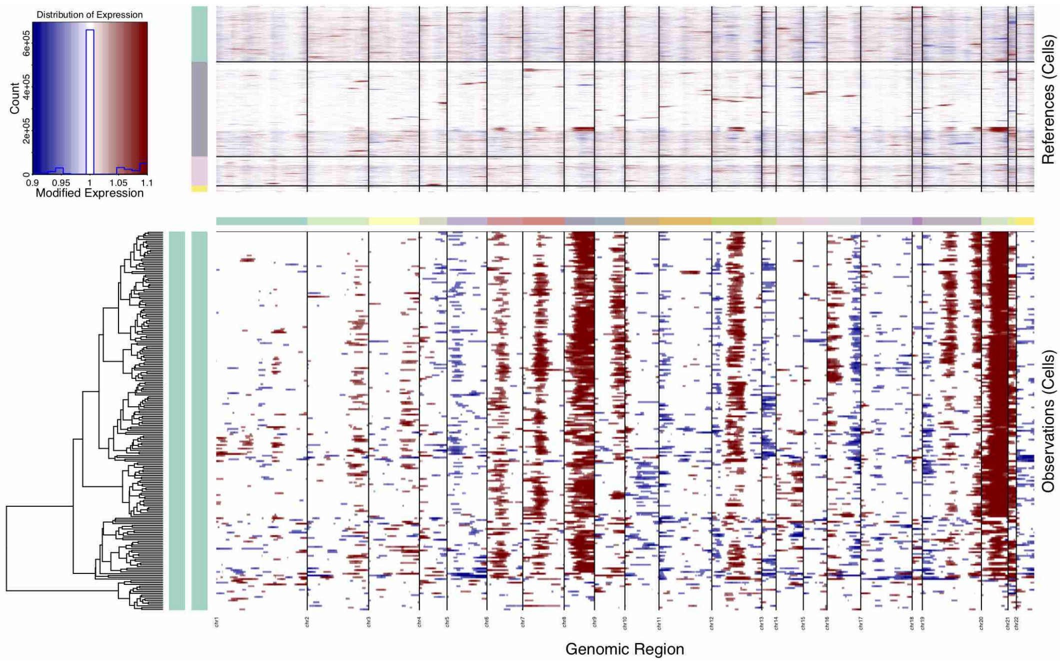

(b) Tumor T76

**Figure S13: Copy number profiles of T59 and T76**. Copy number profiles, inferred from scRNA-seq data using `inferCNV` [14, 15], for cancer epithelial cells of Tumors T59 and T76. Expression values for normal reference cells are plotted in the top heatmap (myeloid cells in green, stromal in grey, lymphocytes in pink, endothelial in yellow). Tumor epithelial cells are plotted in the bottom heatmap, with genes ordered from left to right across the genome. Normal cell expression is effectively subtracted from tumor cell expression, where amplifications are shown in red, and deletions of genomic regions in blue.

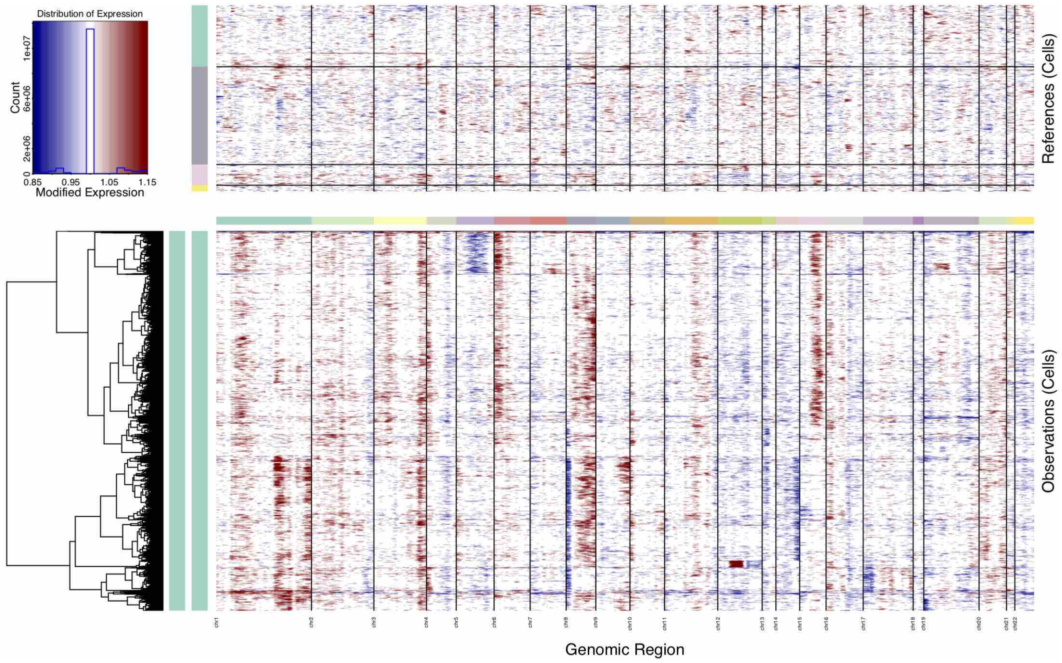

(a) Tumor T77

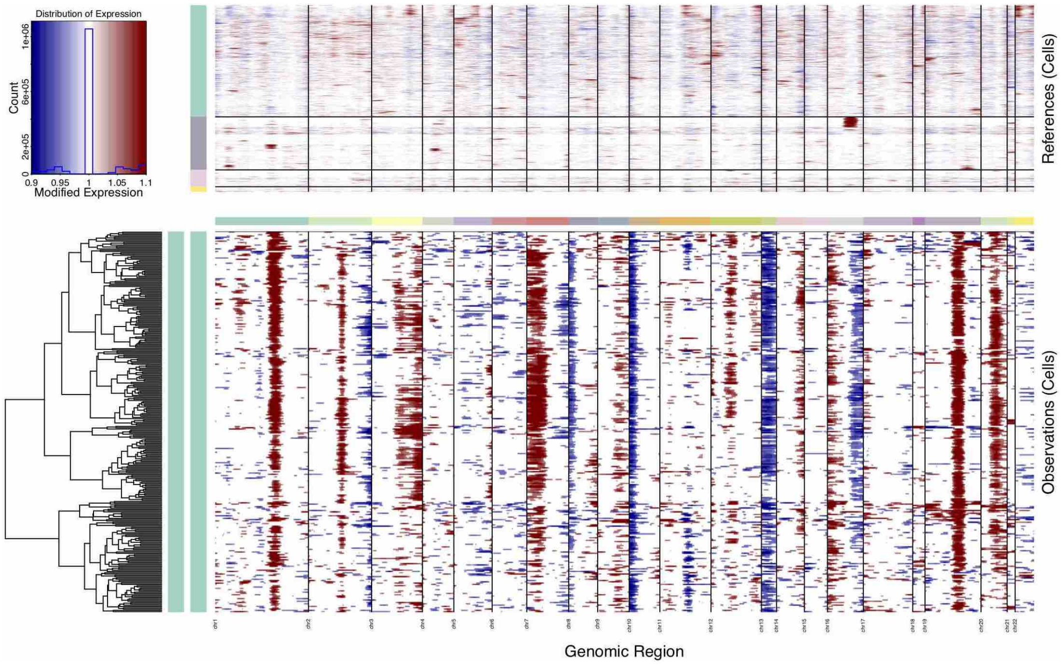

(b) Tumor T89

**Figure S14: Copy number profiles of T77 and T89**. Copy number profiles, inferred from scRNA-seq data using `inferCNV` [14, 15], for cancer epithelial cells of Tumors T77 and T89. Expression values for normal reference cells are plotted in the top heatmap (myeloid cells in green, stromal in grey, lymphocytes in pink, endothelial in yellow). Tumor epithelial cells are plotted in the bottom heatmap, with genes ordered from left to right across the genome. Normal cell expression is effectively subtracted from tumor cell expression, where amplifications are shown in red, and deletions of genomic regions in blue.

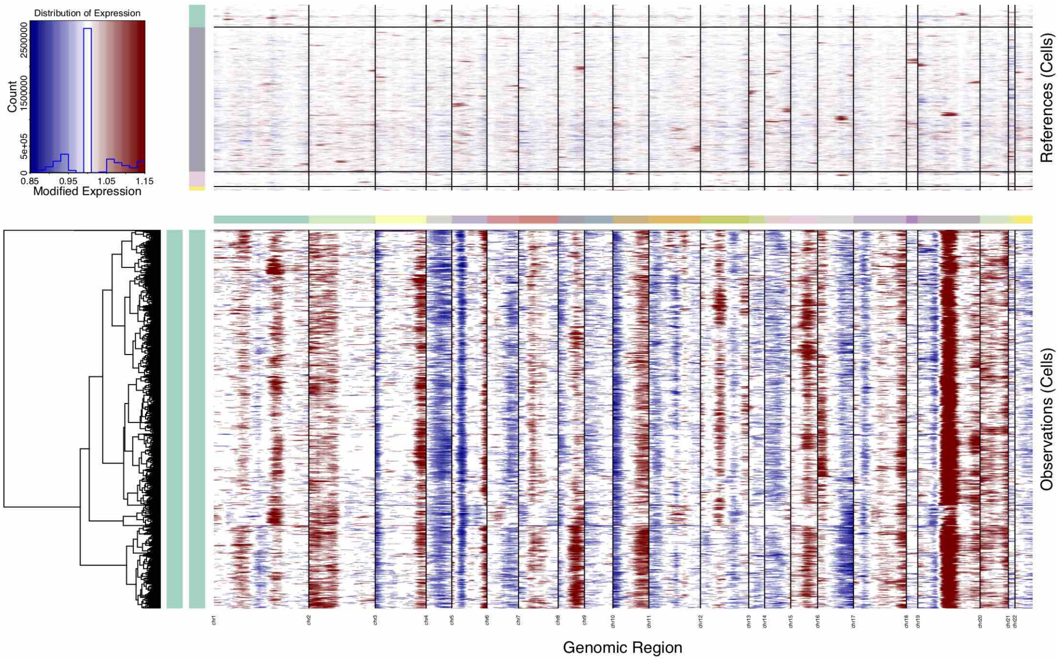

**Figure S15: Copy number profile of T90**. Copy number profiles, inferred from scRNA-seq data using `inferCNV` [14, 15], for cancer epithelial cells of Tumor T90. Expression values for normal reference cells are plotted in the top heatmap (myeloid cells in green, stromal in grey, lymphocytes in pink, endothelial in yellow). Tumor epithelial cells are plotted in the bottom heatmap, with genes ordered from left to right across the genome. Normal cell expression is effectively subtracted from tumor cell expression, where amplifications are shown in red, and deletions of genomic regions in blue.

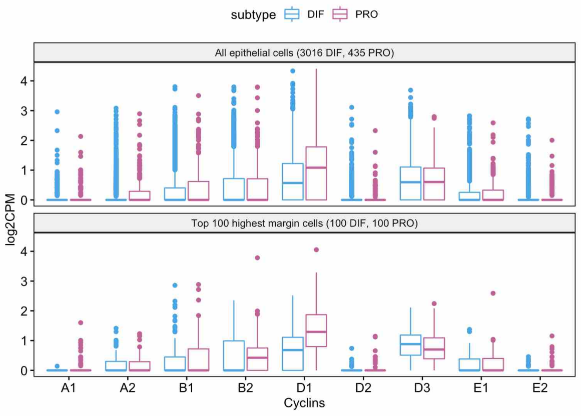

(a) Tumor T59

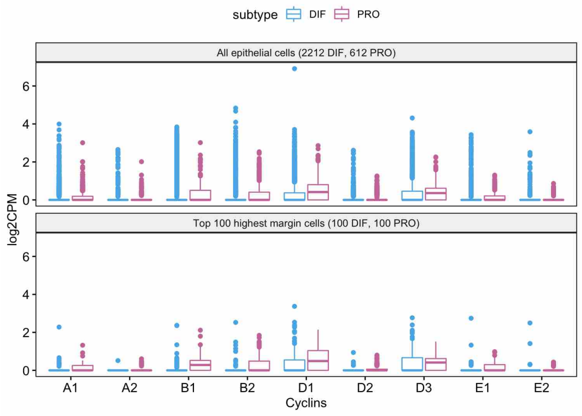

(b) Tumor T77

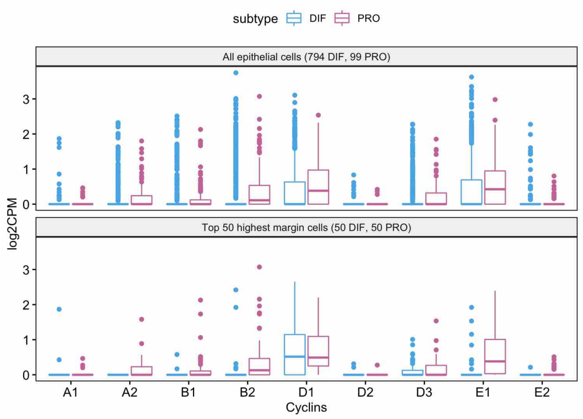

(c) Tumor T90

**Figure S16: Cyclin expression in epithelial cells of tumors T59, T77, and T90**. Shown is the expression level ($y$-axis) of cell cycle phase-specific cyclins ($x$-axis) for epithelial cells of tumors with a substantial proportion of epithelial cells assigned to both the differentiated (blue) and the proliferative subtype (violet). The upper panel shows the comparison of both subtype groups when taking all epithelial cells of a tumor into consideration. The lower panel shows the comparison for a restricted set of epithelial cells that were most confidently classified as either differentiated or proliferative. The restricted view in the lower panel shows an up-regulation of cyclin D1 in proliferative cells of Tumor T59 ($p = 1.8 \cdot 10^{-7}$, Wilcoxon rank-sum test), and of cyclin E1 in proliferative cells of Tumor T90 ($p = 4.9 \cdot 10^{-8}$).

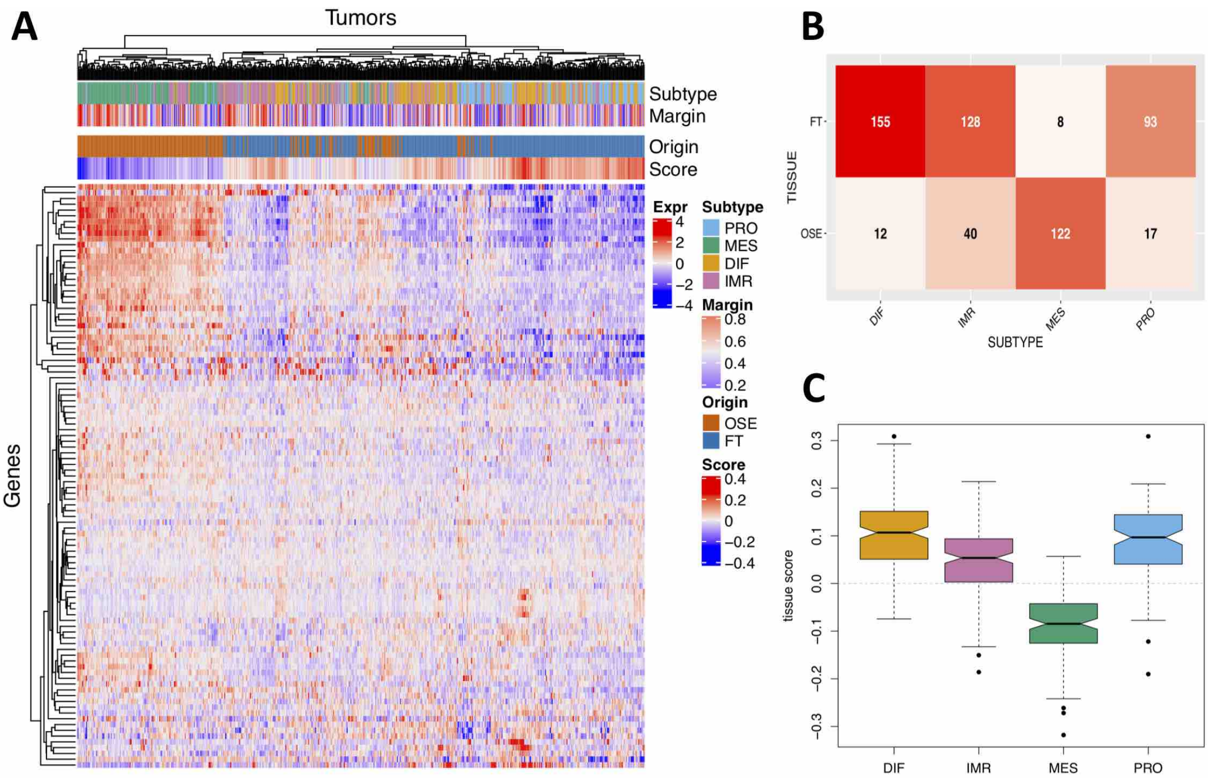

**Figure S17: Tissue of origin.** Plots are shown based on the HGSOC origin signature proposed by Hao *et al.* [31]. The signature consists of 37 genes up-regulated in the fallopian tube (FT) and 75 genes up-regulated in ovarian surface epithelium (OSE). Tissue scores were computed as implemented in the `get.hao.subtypes` function of the `consensusOV` package [19]. Consensus HGSOC transcriptome subtypes were called using the `get.consensus.subtypes` function of the `consensusOV` package. The heatmap in **(A)** depicts centered expression values of the 112-gene signature of Hao *et al.* [31] (rows) across 575 TCGA HGS ovarian tumors (columns). The annotation bars at the top show consensus subtype calls (*Subtype*), associated classification margin scores (*Margin*), predicted tissue of origin (*Origin*), and associated tissue scores (*Score*). The contingency table in **(B)** summarizes the overlaps of the annotation bars *Subtype* and *Origin* from **A**. The table demonstrates that MES calls predominantly coincide with OSE calls, whereas other subtype calls coincide more frequently with FT calls. The boxplot in **(C)** shows tissue scores ($y$-axis, *Score* in **A**) stratified by subtype ($x$-axis, *Subtype* in **A**). The grey dashed line at $y = 0$ indicates the separation of tumors into FT-like expression (score $> 0$) and OSE-like expression (score $< 0$).

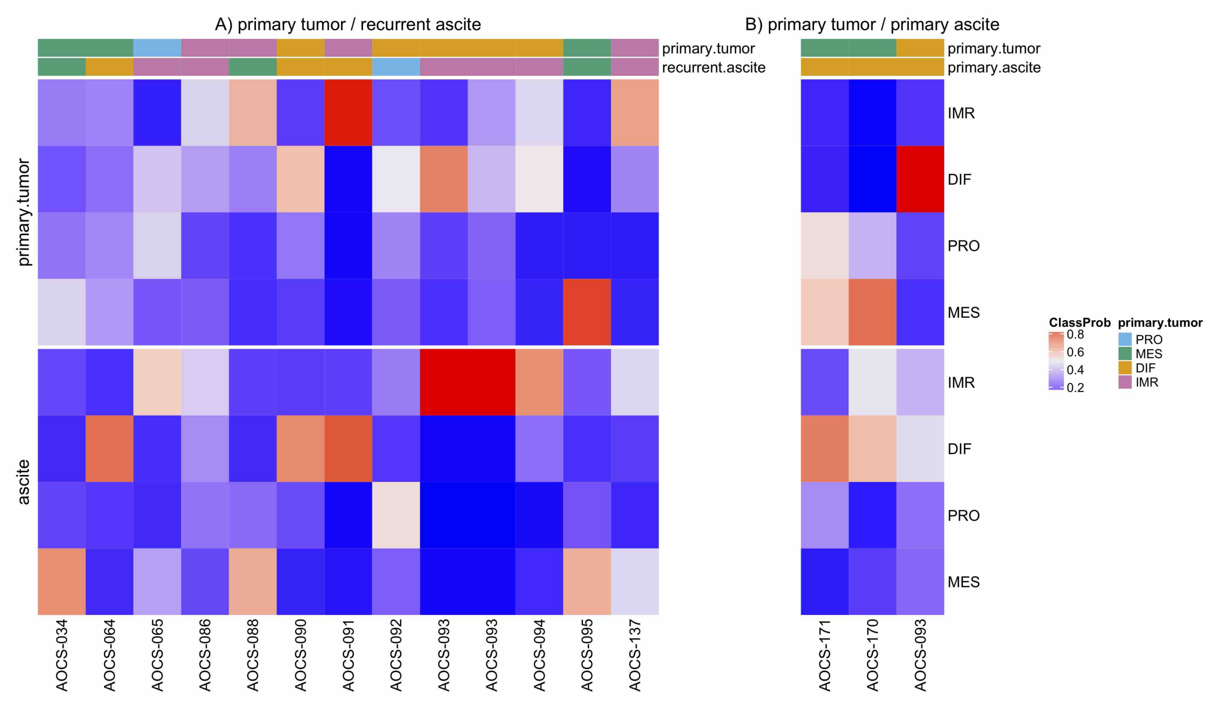

**Figure S18: Subtyping of primary tumor and associated ascites samples.** The heatmap depicts classification probabilities and corresponding subtype calls of the consensus classifier [18] for primary tumor samples and associated **(A)** recurrent and **(B)** primary ascites samples from the HGSOC chemoresistance study of Patch *et al.* [35].

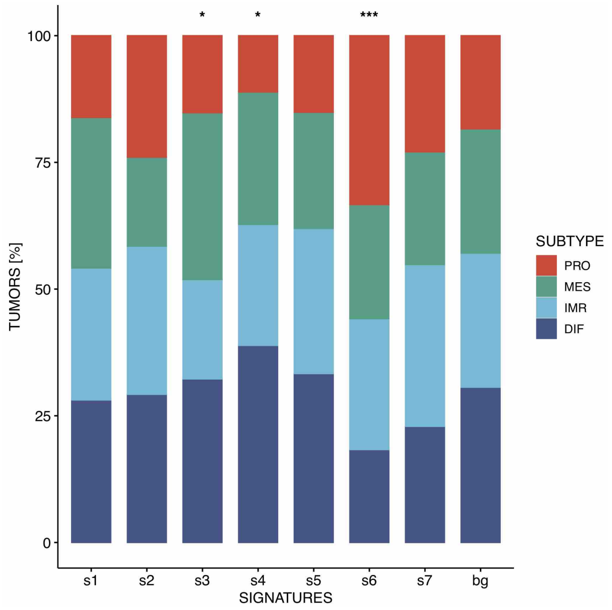

**Figure S19: Subtype association of CN signatures.** The barplot illustrates the relationship between the 7 CN signatures identified by Macintyre *et al.* [32] ($x$-axis) and consensus subtype assignment of 408 TCGA HGS ovarian tumors. Shown is percentage of tumors ($y$-axis), stratified by consensus subtype assignment, exceeding the mean exposure level for each CN signature. The rightmost bar named *bg* shows the background distribution of subtype assignment in all 408 tumors. Significant differences in proportions with respect to the background distribution is indicted by an asterisk ($^{*}p < 0.05$, $^{**}p < 0.01$, $^{***}p < 0.001$, $\chi^2$ goodness-of-fit test, $df = 3$). Number of tumors exceeding the mean exposure level for each signature: 192 ($s1$), 171 ($s2$), 158 ($s3$), 180 ($s4$), 192 ($s5$), 120 ($s6$), 144 ($s7$).
